# Pooled nanoparticle screening using a chemical barcoding approach

**DOI:** 10.1101/2024.09.24.614746

**Authors:** Katherine Vaidya, Michael S. Regan, James Lin, Jenna Houle, Sylwia A. Stopka, Nathalie Y. R. Agar, Paula T. Hammond, Natalie Boehnke

## Abstract

We report the development of a small molecule-based barcoding platform for pooled screening of nanoparticle delivery. Using aryl halide-based tags (halocodes), we achieve high-sensitivity detection via gas chromatography coupled with mass spectrometry or electron capture. This enables barcoding and tracking of nanoparticles with minimal halocode concentrations and without altering their physicochemical properties. To demonstrate the utility of our platform for pooled screening, we synthesized a halocoded library of polylactide-co-glycolide (PLGA) nanoparticles and quantified uptake in ovarian cancer cells in a pooled manner. Our findings correlate with conventional fluorescence-based assays. Additionally, we demonstrate the potential of halocodes for spatial mapping of nanoparticles using mass spectrometry imaging (MSI). Halocoding presents an accessible and modular nanoparticle screening platform capable of quantifying delivery of pooled nanocarrier libraries in a range of biological settings.

---

Pooled screening allows for high-throughput evaluation of nanoparticle (NP) properties that mediate successful delivery outcomes and enable generation of large datasets for subsequent intersystem analyses.^1–8^ Nucleic acid barcodes have been primarily applied to pooled NP screening due to their low detection limits and large number of possible combinations.^9^ They have been used to screen large libraries of lipid nanoparticles (LNPs) to determine tissue accumulation and functional delivery, overcoming limitations associated with optical barcodes such as signal attenuation and spectral overlap.^2,3,5,6,9–21^ However, their large size, degradability, and incompatibility with many nanocarriers, such as those designed for diagnostics and small molecule delivery, limit utility.^22^

There is a need to develop barcoding strategies that are widely applicable to multiple nanocarrier types and expedite targeted therapies for unmet therapeutic needs. For example, ovarian cancer exhibits high mortality rates due to late stage diagnosis and therapy resistance.^23,24^ NPs are a promising alternative drug delivery approach,^25,26^ and pooled screening could accelerate identification of effective therapeutic and diagnostic formulations.

Herein, we report a small molecule barcoding strategy for pooled nanocarrier screening (Scheme 1). We leveraged aryl halide-based barcodes (halocodes) due to their stability, low detection levels,^27,28^ compatibility with both covalent and noncovalent incorporation into nanocarriers, large combinatorial space,^29–32^ and multiplexing potential.

**Scheme 1.**
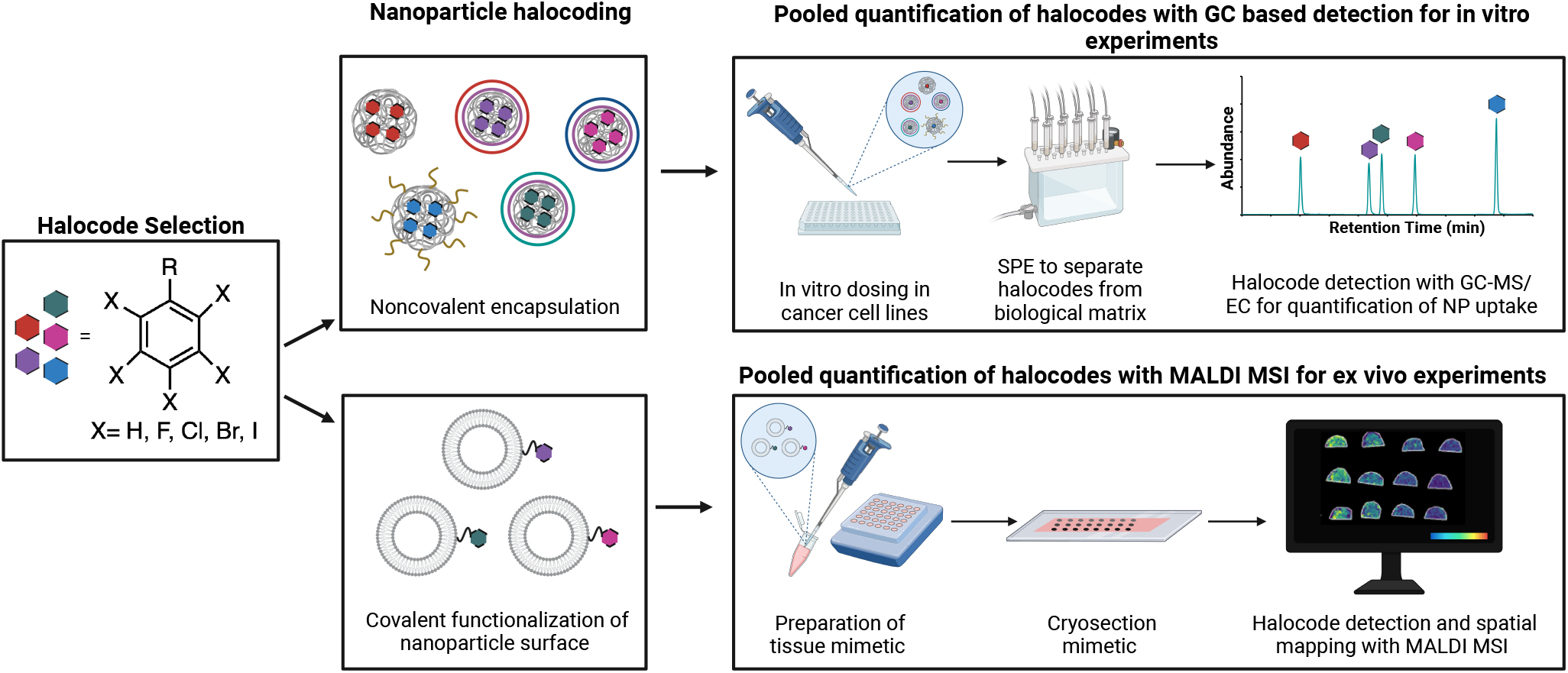
Overview of halocoding platform.

Gas chromatography-electron capture (GC-EC), which identifies analytes based on their ability to capture electrons from a radiation soure, has been the primary mode of quantification for aryl halide tags^27,29,31–33^ due to its sensitive detection of highly electrophilic compounds. GC-mass spectrometry (MS), which detects compounds based on fragmentation patterns after ionization, have also been used in environmental applications.^30,34–38^ While GC-EC can provide more sensitive detection of aryl halides, it is a specialized instrument and less accessible than GC-MS.^39^ Therefore, when developing methods for halocode detection, we initially used a GC with both MS and EC detectors for simultaneous pooled halocode quantification (Figure 1a).

**Figure 1.**
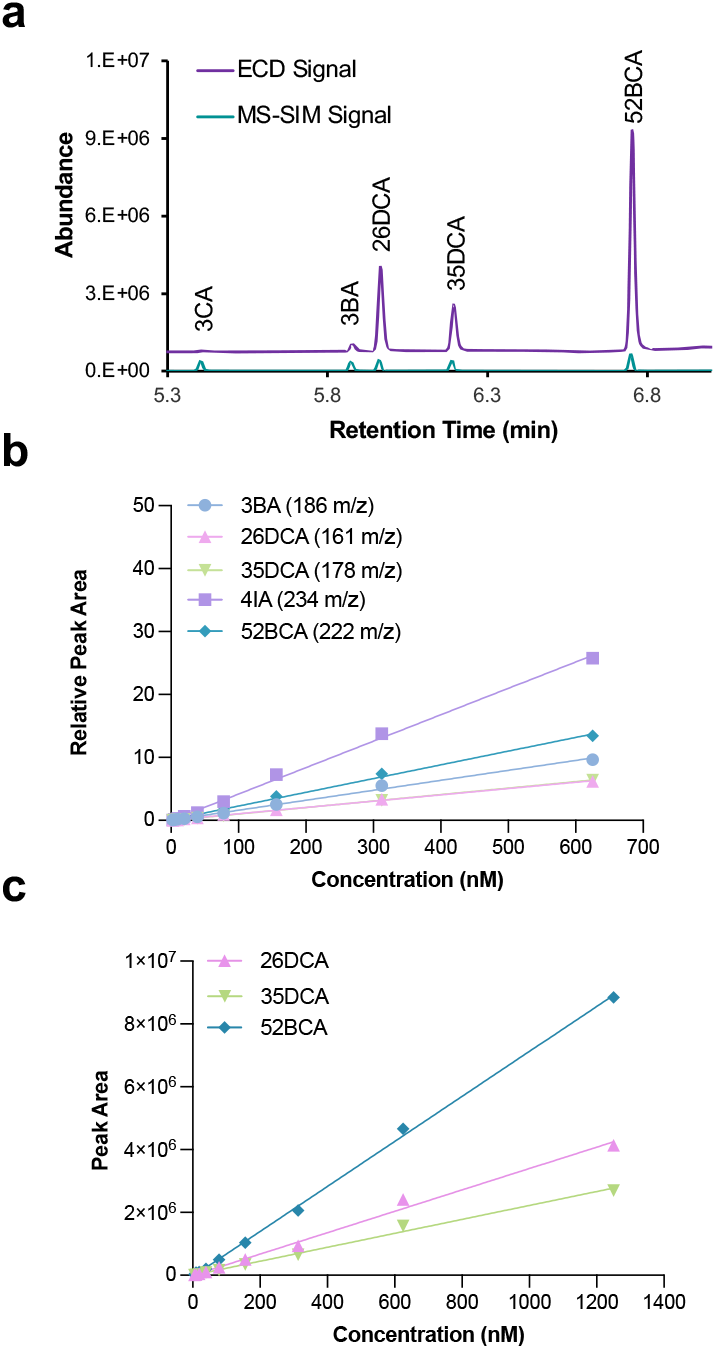
Halocode quantification via GC. (a) GC chromatograms with MS and EC halocode detection. Standard curves of (b) all halocodes with MS and (c) dihalogenated halocodes with EC. 3-bromoanisole (3BA); 3-chloroanisole (3CA); 2,6-dichloroanisole (26DCA); 3,5-dichloroanisole (35DCA); 4-iodoanisole (4IA); 5,2-bromochloroanisole (52BCA).

We observed that all tested halocodes exhibited linear detection with 3-chloroanisole (3CA) as an internal standard using GC-MS, with limits of detection (LODs) below 0.78 nM and limits of quantification (LOQs) below 2.6nM (Figure 1b and Table S1). Conversely, only halocodes with multiple halide substituents showed linear response with GC-EC at low concentrations (Figure 1c), highlighting a known limitation of this technique-that sensitivity of EC-based detection can vary greatly with analyte structure.^40^ Although factors including ionization efficiency can cause variation in MS signal intensities, this effect is comparatively minor.^41^ We elected to use GC-MS for halocode quantification in this study as we reasoned barcodes should have similar signal intensities to enable detection of pooled barcodes via a single GC method.

We synthesized a set of five nanoparticle formulations, each with a unique halocode encapsulated inside (Figure 2a). While below the possible number of halocode combinations that can be used in a single experiment, we chose to focus on a limited number of barcodes to optimize several key steps of our halocoding platform, including GC method, halocode encapsulation, and extraction from biological matrices. Additionally, simultaneous detection of five analytes can be challenging through traditional fluorescence detection, demonstrating that even small halocoded pools can be used to gain valuable insights into nanoparticle delivery. The following aryl halides were used as halocodes: 4-iodoanisole (4IA), 5-bromo-2-chloroanisole (52BCA), 2,6-dichloroanisole (26DCA), 3-bromoanisole (3BA), and 3,5-dichloroanisole (35DCA).

**Figure 2.**
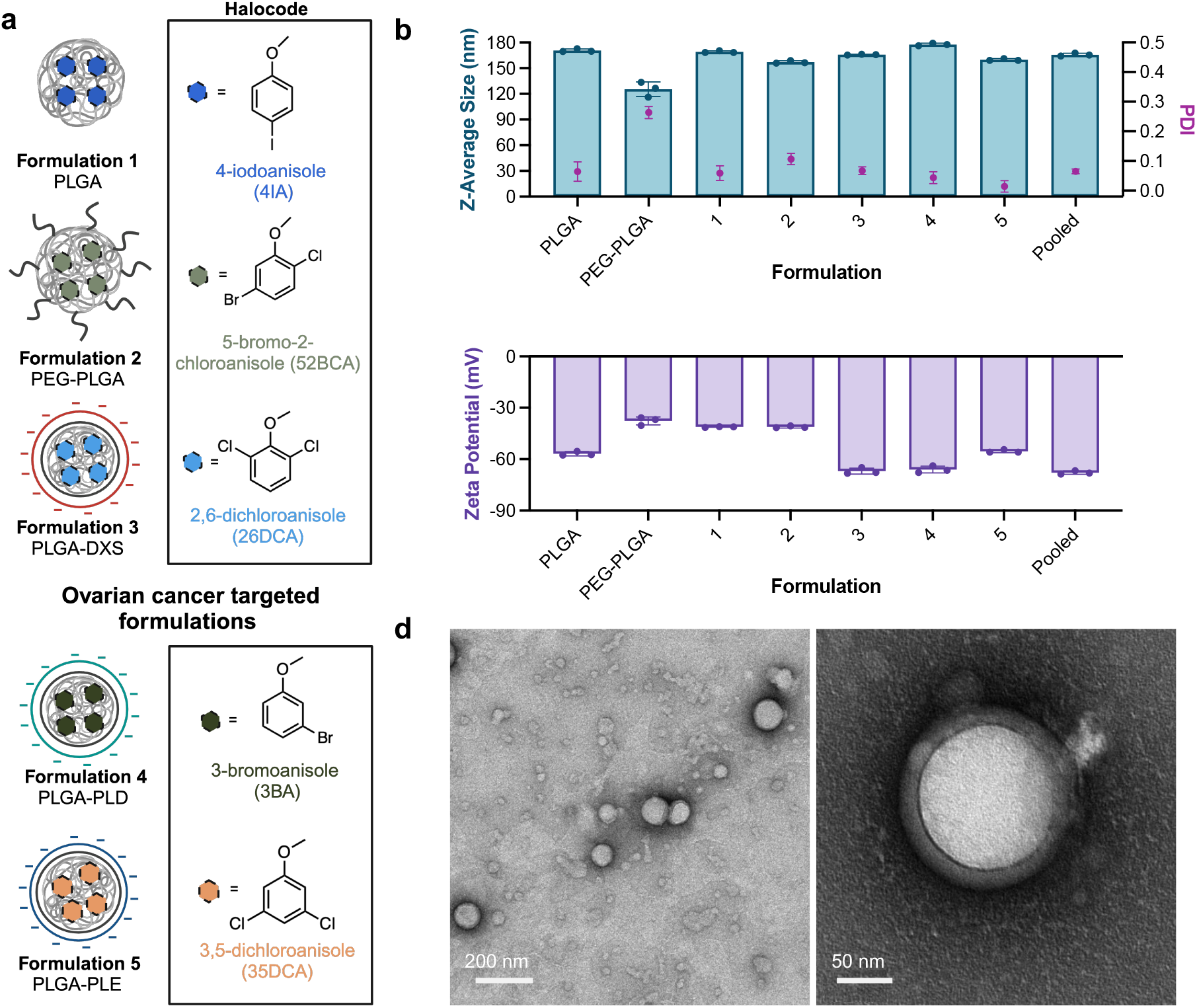
Halocoded nanoparticle library design and characterization. (a) Halocoded PLGA NP library. (b) Hydrodynamic size and PDI of the NP library measured by DLS. (c) Zeta potential of the NP library in milliQ H_2_O measured by electrophoretic light scattering. Data shown as mean ± SD of three repeat measurements. (d) Representative bright-field transmission electron micrographs of the pooled nanoparticle library. NPs were negatively stained with 2% uranyl acetate.

We chose anisole-based halocodes due to their hydrophobicity for loading into poly(D,L-lactide-co-glycolide) (PLGA) NPs via nanoprecipitation.^42,43^ Anionic PLGA-based formulations were chosen due to their biocompatibility, degradability, and ease of synthesis.^44^ We evaluated the effects of solvent, polymer concentration, and ratio of halocode to polymer on halocode encapsulation. Our primary goal was to minimize overall halocode loading, reducing any potential impact of halocodes on NP properties. We achieved halocode loading levels ranging from 0.35-0.64 wt. % (Figure S3 and Table S2), which correspond to halocode concentrations well above determined LOQ values. This is particularly useful for quantification of untargeted nanocarriers, which can have low and variable cellular uptake.^45,46^ Moreover, using halocode loading levels lower than other reporter molecules, including fluorescent dyes, which can range from 1-30 wt. % in analogous systems, and nucleic acid based barcodes^2,47,48^ can potentially mitigate challenges associated with high probe loading, such as altered NP properties and functional delivery profiles.^4,8,,47–49^

We used electrostatic layer-by-layer (LbL) assembly to modify PLGA nanoparticles with unique surface chemistries. Poly-L-arginine (PLR) was deposited as a common first layer followed by deposition of dextran sulfate (DXS), poly-L-aspartate (PLD), or poly-L-glutamate (PLE) to provide a range of targeting properties in the context of ovarian cancer.^23,50,51^ Additionally, a polyethylene glycol (PEG)-PLGA formulation was included due to the clinical utility of PEG as an antifouling polymer.^52^ We hypothesized that these NPs would yield diverse uptake patterns, and the ability to confirm these trends with halocode screening could demonstrate the accuracy and utility of our platform.

Dynamic light scattering (DLS) and transmission electron microscopy (TEM) were used to characterize the resulting NPs (Figure 2b-d, Figure S1). We targeted NP diameters less than 200 nm, a size compatible with cellular uptake.^53^ All formulations had a narrow size distribution with a polydispersity index (PDI) of less than 0.2^54^ and a zeta potential of less than -30 mV, which is sufficient to maintain colloidal stability.^50^ We confirmed that halocode encapsulation did not significantly change NP diameter, dispersity, or zeta potential (Figures 2b-d, S2). The halocoded library also remained stable when pooled. Additionally, no aggregation or changes in particle morphology were observed when imaging pooled NPs via TEM (Figure 2d).

To determine if halocoding impacts nanoparticle behavior in vitro, we separately loaded three unique halocodes (3CA, 26DCA, and 35DCA) into PLGA NPs along with a fluorophore (Cy5) for downstream fluorescence-based analysis (Figure S4). NP-cell association of halocoded Cy5-NPs was assessed using flow cytometry following 24 hr incubation with the high grade serous ovarian cancer cell line OVCAR8, which was selected due to the need for targeted therapies in ovarian cancer and extensive prior characterization.^2,23,50,55^ We observed cells efficiently took up both halocoded and non-halocoded PLGA nanoparticles with minimal differences in fluorescence intensity (Figure S5a). Cells were also imaged via fluorescence microscopy, and we observed that both halocoded and non-halocoded PLGA nanoparticles were efficiently and similarly internalized in vesicles within the cytosol (Figure S5b). Additionally, while aryl halides vary in biocompatibility, many are deemed nontoxic per their safety data sheets (SDS). We conducted viability tests and observed no dose-dependent toxicity for four representative halocodes up to at least 0.25 mM, exceeding quantitation limits by over 250,000-fold (Figure S6).

We used reversed-phase solid phase extraction (SPE) to isolate halocodes from cell suspensions for subsequent GC analysis.^56,57^ We were initially able to achieve 84±4.5% average recovery for halocodes spiked into cell lysate (Figure S7a). However, when increasing the density of cells in the sample matrix from 0.2e6 to 2.0e6 cells, halocode extraction efficiency decreased to 71±14% and became more variable depending on halocode polarity (Figure S7b). Following SPE procedure, we achieved 96±13% recovery for spiked halocodes and 88±5.4% for halocoded NPs from the concentrated cell lysate matrix (Figure S7c,d).

Each of the formulations from the halocoded NP library was incubated with OVCAR8 cells for four hours (Scheme 1, Figure 3a).^2,23,45^ As halocodes are non-covalently incorporated into the tested PLGA NPs, we confirmed that potential premature halocode release would not confound our studies using 52BCA (Figure S8). Following pooled SPE, we found that cell association of PLGA-PLD and PLGA-PLE NPs were 29±2.8% and 20±4.4%, respectively, which were higher than the cell association of the PLGA, PEG-PLGA, and PLGA-DXS formulations, which ranged from 3.3±0.3% to 3.9±1.6% (Figure 3b). This is in line with our prior findings that NPs layered with PLD and PLE preferentially target ovarian cancer cells.^2,23,25,50,55,58^

**Figure 3.**
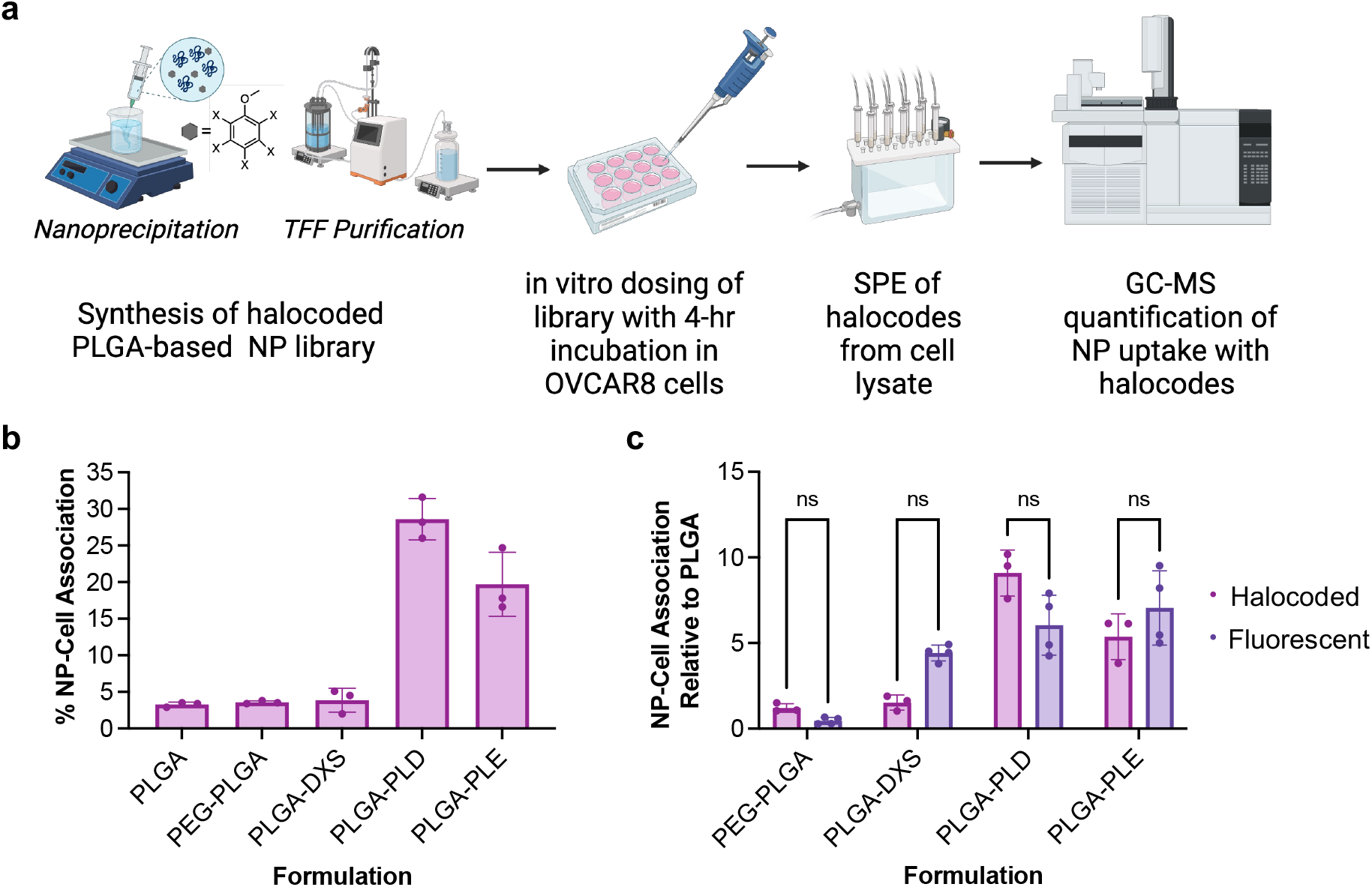
Pooled quantification of halocoded nanoparticle-cell association in vitro. (a) Schematic of the experimental workflow for in vitro dosing of the halocoded NP library. (b) NP-cell association of the halocoded NP library as a function of the administered NP maximum dose (n=3) (c) Comparison of the NP-cell association of NP library formulations relative to bare PLGA with halocode quantification via GC-MS (n=3) and Cy5 fluorescence quantification via microplate spectrophotometry (n=4). Data shown as mean ± SD.

To directly compare results from halocode quantification to more conventional and unpooled fluorescence-based detection, we evaluated the NP library in an unpooled manner with encapsulated Cy5 instead of halocodes (Figure 3c, S9). We determined no significant differences in the relative cell association between the two libraries, indicating the suitability of halocode detection for in vitro NP screening experiments.

While GC-MS provides a rapid method for quantifying diverse molecules, we sought to establish the use of matrix assisted laser desorption ionization (MALDI) mass spectrometry imaging (MSI) to provide both quantitative and spatial information of halocoded NPs.^59^ As MSI can be used to visualize the spatial distribution of molecules in a biological sample by measuring corresponding mass to charge ratios (*m/z*),^60^ MSI-based detection of halocoded NPs could enable spatial quantification of NP accumulation with the potential to multiplex with omics techniques, such as lipidomics and proteomics.^61^

We elected to use liposomes for their clinical utility and the ease with which lipids can be detected via MALDI-MSI.^62^ Liposomes were formulated to include a propargyl containing phospholipid with a phosphatidylethanolamine head-group (18:0 PE) as a reactive handle for covalent halocoding via copper catalyzed azide-alkyne cycloaddition, selected due to its high efficiency and compatibility with aqueous colloidal systems.^55,63^ We selected three representative halocodes, 3-chlorophenol (3CP), 3,5-dichlorophenol (35DCP), and 2,3,6-trichlorophenol (236TCP), and modified them with an ethylene glycol linker bearing a reactive azide end group^29^ (Figures S10-S15) to yield 3-chlorophenol-EG-N_3_, 3,5-dichlorophenol-EG-N_3_, and 2,3,6-trichlorophenol-EG-N_3_, respectively. Propargyl liposomes were separately halocoded^55^ following thin-film hydration (Figures 4a, S16).

**Figure 4.**
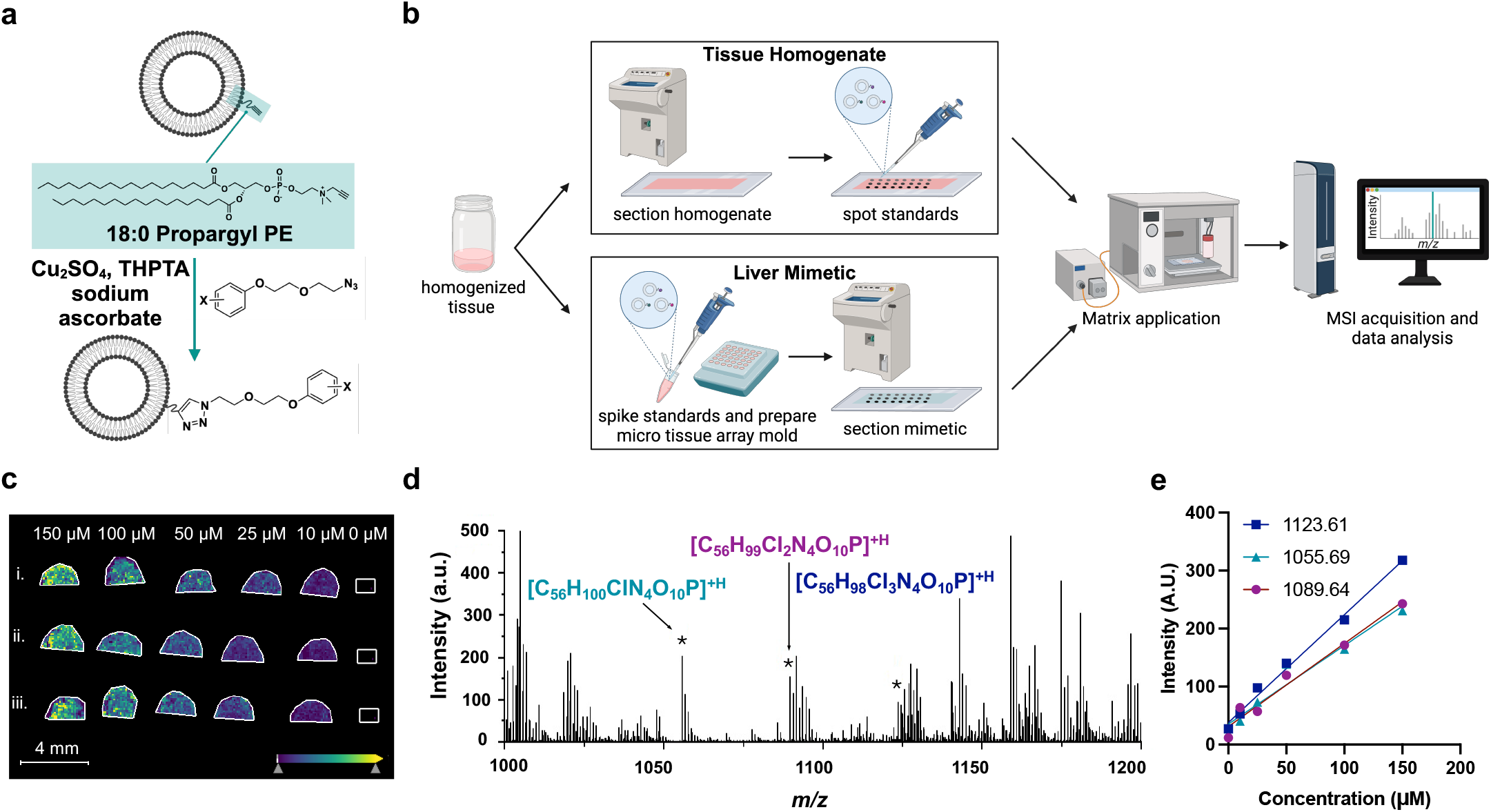
MALDI MSI for detection of covalently halocoded liposomes. (a) Covalent halocoding via copper catalyzed alkyne-azide cycloaddition. (b) Overview of sample preparation workflows. (c) MS signal intensities of halocoded lipids spotted on tissue homogenate. Integrated signal intensities corresponding to *m/z* values of i) 1055.7 (3CP-EG-lipid); ii) 1089.6 (35DCP-EG-lipid); iii) 1123.6 (236TCP-EG-lipid) are shown. (d) Representative mass spectra of halocoded lipids in tissue homogenate. (e) Signal-response calibration curves of integrated signal intensities of halocoded lipids incorporated into tissue mimetic.

We first deposited standard solutions of halocoded liposomes, separately and in pooled format, onto a murine tissue homogenate (Figure 4b, Figure S17). We observed specific mass to charge ratios corresponding to each halocoded lipid in a concentration-dependent manner. Moreover, signal for each lipid was also detected in the pooled sample, demonstrating the specificity of this method for detecting individual analytes in a complex mixture.

Next, the halocoded liposomes were incorporated into a murine liver mimetic where homogenized liver tissue was spiked with halocoded liposomes and frozen into a support mold to simulate distribution of liposomes throughout tissue (Figure 4b,d). We elected to use a liver mimetic as the liver is the main clearance organ of liposomes.^64^ Integration of signal intensities again revealed a concentration-dependent and linear relationship down to low micromolar concentrations (Figures 4e, S18). As greater than 50% of an injected liposome dose tends to accumulate in the liver ^65^, we believe this detection range to be useful for in future vivo quantification.

In conclusion, we report the development of a NP barcoding technology using aryl halide-based tags (halocodes) for pooled NP screening. We used halocoding for pooled GC-MS quantification of a PLGA-based NP library to identify formulations that preferentially target ovarian cancer cells and verified the accuracy of the trends in NP-cell association with conventional unpooled fluorescence-based quantitation methods. We also applied halocoding for quantification and spatial mapping of liposomes using MALDI MSI. Given compatibility of halocodes with multiple classes of nanocarriers, modularity of halocode attachment strategies, extensive halocode parameter space, and low limits of quantitation of halocodes with GC-MS/EC and MSI technologies, we anticipate that halocoding will serve as a valuable barcoding tool to expedite NP screening for the broader scientific community.

## ASSOCIATED CONTENT

### Supporting Information

The Supporting Information is available free of charge on the ACS Publications website.

Details about materials and methods, supplemental figures and tables (PDF).

## AUTHOR INFORMATION

### Author Contributions

K. V.: experimental, formal analysis, writing – original draft, review and editing. M. S. R.: experimental, formal analysis, writing – review and editing. J. L.: experimental, writing – review and editing. J. H.: experimental, writing – review and editing. S. A. S.: experimental, formal analysis, writing – review and editing. N. Y. R. A.: formal analysis, writing – review and editing. P. T. H.: formal analysis, writing – review and editing. N. B.: conceptualization, experimental, formal analysis, writing – original draft, review and editing, funding acquisition. All authors have given approval to the final version of the manuscript.

## ACKNOWLEDGMENT

This work was supported by the National Cancer Institute (NCI) (K99CA255844, 4R00CA255844). Mass spectrometry imaging work was supported by the National Institute of Biomedical Imaging and Bioengineering (P41EB028741 supporting N.Y.R.A). This work was also supported in parts by the Koch Institute (core) Grant P30-CA14051 from the National Cancer Institute and the University of Minnesota Characterization Facility, which receives partial support from the NSF through the MRSEC (DMR-2011401) and the NNCI (ECCS-2025124) program. We thank A. N. Koehler, P. A. Clemons, and S. L. Schreiber for thoughtful discussions surrounding this work.

