## Supporting information, figures, and tables for "Pooled nanoparticle screening using a chemical barcoding approach"

**Contents**

Materials S2

Methods S3

Supplemental Figures S11

Supplemental Tables S25

References S26

### Materials

*Reagents*

Poly(D,L-lactide-co-glycolide) (Resomer 502H, 7 kDa: 17 kDa, acid terminated), Poly(D,L-lactide-co-glycolide) 50:50-b-PEG (10 kDa PLGA, 2kDa PEG), 3-chloroanisole, 3-bromoanisole , 3,5-dichloroanisole, 5-bromo-2-chloroanisole, 4-iodoanisole, dextran sulfate salt, acetone, HPLC grade acetonitrile, HPLC grade methanol, and HPLC grade DMSO were purchased from Sigma. Cy5 free acid dye was purchased from Lumiprobe. Poly-L-arginine hydrochloride(PLR200, 38.5 kDa), poly-L-aspartic acid sodium salt (PLD100, 14 kDa), and poly-D,Lglutamic acid sodium salt (PRE100, 15 kDa) were purchased from Alamanda Polymers. Lipids and cholesterol were purchased from Avanti Polar Lipids. Bromo-PEG1-azide was purchased from BroadPharm. All other reagents were acquired from Sigma. Reagents and instrumentation used for MALDI-MSI are described in the text.

Conical tubes and borosilicate glass scintillation vials were purchased from VWR. 1.7mL Eppendorf tubes were purchased from Genesee Scientific. 1cc 28Gx1/2” insulin syringes were purchased from Amazon. D02-E100-05-N and C02-E100-05-N tangential flow filtration filters and ACTU-P13-25N PharmaPure #13 Tubing were purchased from Repligen. Polystyrene semi-micro cuvettes for the Malvern Zetasizer were purchased from Sarstedt and folded capillary cells were purchased from Fisher Scientific. Carbon coated mesh 400 copper grids were purchased from Electron Microscopy Sciences. 2% uranyl acetate (UA) solution was purchased directly from the UMN Characterization Facility.

85% phosphoric acid was purchased from Fischer Chemical. Oasis 1cc HLB and 3cc HLB PRiME SPE cartridges were purchased from Waters. Clear 2mL screw top vials, 400µL flat bottom glass inserts, PTFE/red silicone screw caps, and all other GC consumables were purchased from Agilent.

RPMI-1640 (ATCC), 1x PBS solution pH 7.4 (Corning), heat inactivated fetal bovine serum (Gibco) , trypsin EDTA (Corning), and penicillin streptomycin (Corning) were purchased from Fisher Scientific. Tissue culture plasticware was purchased from Genesee Scientific. 10x RIPA lysis buffer was purchased from Neta-Scientific. Sodium dodecyl sulfate (SDS) was purchased from Sigma.

*Cells*

The OVCAR8 cells were a gift from the Hammond Lab. Cells were cultured at 37°C with 5% $CO_2$. Cell line-specific culture information is provided in the ‘In Vitro Dosing of Halocoded NP Library’ and ‘In Vitro Dosing of Fluorescent NP Library’ sections.

### Methods

The nanoparticle related methods include details for synthesis, purification, and quantification of the encapsulated halocodes. The methods have been partially adapted from references Boehnke et al.[1,2], Kong et al.[3] , and Shin et al.[4].

*Synthesis of PLGA Nanoparticles*

PLGA was dissolved at a concentration of 10 mg/mL in acetone. 3 mL of milliQ water were added to a 20 mL scintillation vial and stirred at 500 rpm. 500 µL of PLGA solution was drawn into a syringe with a 28.5-gauge needed attached. In the scintillation vial, the tip of the needle was submerged below the water line and PLGA solution was slowly injected into the water under constant stirring. An additional 7 mL of milliQ water was added to the solution to dilute nanoparticle concentration to 0.5 mg/mL. The solution was left to stir for 3 hours to allow for complete evaporation of the solvent.

*Synthesis of PEG-PLGA Nanoparticles*

PEG-PLGA (50:50, 10k PLGA, 2k PEG) was dissolved at a concentration of 10 mg/mL in a 1:1 solution of DMSO: acetone. 3 mL of milliQ water were added to a 20 mL scintillation vial and stirred at 500 rpm. 500 µL of PEG-PLGA solution was drawn into a syringe with a 28.5-gauge needle attached. In the scintillation vial, the tip of the needle was submerged below the water line and PEG-PLGA solution was slowly injected into the water under constant stirring. An additional 7 mL of milliQ water was added to the solution to dilute nanoparticle concentration to 0.5 mg/mL. The solution was left to stir for 3 hours to allow for complete evaporation of the solvent.

*Synthesis of Halocoded PLGA and PEG-PLGA Nanoparticles*

PLGA was dissolved at a concentration of 50 mg/mL in acetone and the aryl halide (AH) of choice (3BA, 26DCA, 35DCA, or 4IA) was dissolved at a concentration of 50 mg/mL in acetonitrile for PLGA nanoparticles. PEG-PLGA was dissolved at a concentration of 50 mg/mL in a 1:1 solution of DMSO: acetone and heated for 15 min at 65°C for complete solubilization of the polymer. 52BCA was dissolved at a concentration of 50 mg/mL in DMSO for PEG-PLGA nanoparticles. 3 mL of milliQ water were added to a 20 mL scintillation vial and stirred at 500 rpm. For 3BA, 26DCA, 35DCA, and 52BCA halocoded NPs, the halocode was loaded at 25 wt.% with respect to the NP core. For 4IA halocoded NPs, the halocode was loaded at 10 wt.% with respect to the NP core. In a 2mL Eppendorf tube, 50 µL or 20uL of AH solution, 200 µL of polymer solution, and 350 µL of acetone were mixed and drawn into a syringe 28.5-gauge needle attached. In a scintillation vial, the tip of the needle was submerged below the water line and the PLGA-AH or PEG-PLGA-AH solution was injected into the water under constant stirring. An additional 13 mL of milliQ water was added to the solution to dilute the nanoparticle concentration to 0.5 mg/mL. The scintillation vial was capped and left to stir for 10 min before purification by tangential flow filtration as described below.

*Synthesis of Fluorescent PLGA and PEG-PLGA Nanoparticles*

PLGA was dissolved at a concentration of 10 mg/mL in acetone and Cy5 free acid dye was dissolved at a concentration of 50 mg/mL in DMSO for PLGA nanoparticles. PEG-PLGA was dissolved at a concentration of 50 mg/mL in a 1:1 solution of DMSO: acetone and heated for 15 min at 65°C for complete solubilization of the polymer. 6 mL of milliQ water were added to a 20 mL scintillation vial and stirred at 500 rpm. In a 2mL Eppendorf tube, 2μL of dye was added to 1 mL of polymer solution, mixed and drawn into a syringe 28.5-gauge needle attached. In a scintillation vial, the tip of the needle was submerged below the water line and the polymer-fluorophore solution was injected into the water under constant stirring. An additional 2 mL of milliQ water was added to the solution prior to purification using tangential flow filtration as described below.

*Tangential Flow Filtration (TFF)*[5]

To remove unencapsulated AHs after nanoparticle synthesis or to purify nanoparticles after layering, nanoparticle solution in a conical tube was connected to a Repligen KrosFlo KR2i TFF system via Masterflex Teflon coated PharmaPure tubing. D02-E100-05-N membranes were used to purify particles. Samples were run at flow rates of 13 mL/min with size 13 tubing with milliQ water as the exchange buffer. Samples were washed until five volume equivalents were collected in the permeate and then recovered by reversing the direction of the peristaltic pump and backflushing with 1 mL of milliQ water. Once purified, samples were concentrated with C02-E100-05-N membranes and recovered using the same method. Following TFF, nanoparticles were characterized dynamic light scattering.

*Layer-by-Layer Functionalization of Nanoparticles*

PLGA nanoparticles were layered by adding equal volumes of nanoparticle solution at 1.0 mg/mL to polyelectrolyte solution under sonication at room temperature. The mixture was sonicated for around 3 seconds. The optimal weight equivalent (wt. eq.) for each layer was determined through a polyelectrolyte titration using 50 μL samples of the nanoparticles for each tested wt. eq. Each test ratio was layered as described above and then characterized. If the resulting nanoparticles had a zeta potential with a magnitude greater than 30 mV and a size within 10 nm of the size of the bare nanoparticles, it was chosen as the optimal wt. eq. for that layer. The wt. eqs. of the cationic first layer, PLR, were 0.2 for the PLGA nanoparticles. The wt. eq. for each anionic polyelectrolyte layer were: 0.6 wt. eq. PLD, 0.6 wt. eq. PLE, and 0.6 wt. eq. DXS. All polyelectrolyte solutions were prepared in water. Layering was also carried out in water. The layered particles were purified via TFF as described above and characterized (see 'DLS Characterization of Nanoparticles' and 'TEM Imaging of Halocoded Nanoparticles').

*DLS Characterization of Nanoparticles*

Nanoparticle hydrodynamic size and polydispersity were measured using dynamic light scattering (Malvern Zetasizer Pro, λ = 633 nm). Zeta measurements were acquired with the Malvern Zetasizer Pro using laser doppler electrophoresis. Nanoparticle solutions were diluted in milliQ water in polystyrene cuvettes for size measurements and DTS1070 folded capillary cuvettes for zeta measurements.

*HPLC for Quantification of Halocode Loading*

HPLC characterization was carried out on an Agilent 1260 Infinity II system with a G711B quaternary pump, G7129A autosampler, G7116A MCT, and a G715A DAD WR. The nanoparticles were dissolved in a volume of 3 parts ACN to 1 part NP suspension to release the encapsulated halocodes, and a Poroshell 120 EC-C18 column (4.6 x 100 mm, 2.7 µm) was utilized to analyze the weight % loading of halocodes with respect to PLGA nanoparticle core using a gradient of 50-95% ACN/$H_2O$ over 8.1 minutes with a flow rate of 1.5 mL/min. The halocodes eluted in a time range from approximately 3.2- 5.2 minutes in order of decreasing polarity.

*TEM imaging of Halocoded Nanoparticles*

Images of barcoded nanoparticles were acquired using a FEI Tecnai T12 TEM. Carbon coated mesh 400 copper grids were glow discharged and 10 µL of PLGA nanoparticles at a concentration of 0.25 mg/mL were deposited on the grid. The grid was blotted after 1 minute to remove excess sample. 10 µL of 2% uranyl acetate (UA) solution was deposited on the grid and immediately blotted. After 30 seconds the UA staining process was repeated, and the grid was left to dry for 10 minutes. The grid was mounted on a single tilt sample holder. The microscope was operated at 120 kV with a magnification of 10,000-40,000x for assessing particle morphology and size distribution. All bright-field images were recorded on a Gatan MSC794 CCD Camera

*GC-MS/EC Detection of Halocodes*

An HP-5MS IU (5%-phenyl)methylolysiloxane column (30m x 0.250mm x 0.25um, Agilent) was used during halocode analysis on an Agilent 8890 GC coupled with an Agilent 5977B MS and Agilent G2397A ECD. The two detectors were connected to the column with 0.15mm diameter restrictor tubing using a three-way splitter plate with half of the column effluent flowing to each detector. The helium carrier gas flow was set at 1.2 mL/min respectively and the 5% methane/ 95% argon make-up gas flow for the ECD was set at 30mL/min. Standards containing increasing concentration of halocodes with fixed levels of an aryl anisole internal standard, and eluent from SPE in either methanol or acetonitrile were injected at a volume of 1µL in the splitless mode (30sec delay), with the following oven temperature program: 45°C(1min), ramp to 250°C at 30°C/min and hold at 250°C for 0.2min. Quantification of the halocodes was conducted under selected ion monitoring mode (SIM) with MSD.

*SPE of Halocoded Nanoparticles with 1cc Oasis HLB Cartridge for Matrix Recovery Studies*

Halocoded nanoparticles at a concentration of 0.5 mg/mL in milliQ water were added to either water, PBS, full biological media used for cell culture (RPMI-1640 media + 10% FBS + 1% penstrep), or cell lysate. The nanoparticle samples were diluted at a 1:1 ratio with 4% phosphoric acid to acidify samples before SPE with a 20-position cartridge vacuum manifold. The Oasis HLB 1cc SPE cartridge was conditioned with 2 mL of methanol and equilibrated with 1 mL of milliQ water. The sample was loaded at a flow rate of 1 mL/min, washed with 1 mL of water and 1 mL of 5% methanol, and then eluted with 1 mL of methanol at 1 mL/min. The halocode concentrations of eluted sample were determined with GC-MS and compared to the initial halocode concentration calculated based on the wt. % loading of the nanoparticle to determine the SPE efficiency for each halocode. In general, recovery ranges of analytes with optimized SPE procedures can range between 70-120%, which can account for loss of analyte during the extraction process or enhanced analyte signal from minor matrix effects.[6–8]

*Optimization of SPE Protocol for Extraction of Halocoded Nanoparticles from Cell Lysate*
We elected to focus SPE optimization studies on cell lysate as the matrix, to mimic in vitro conditions. We chose to use 2.0e6 cells in these optimization studies as this is the expected maximum number of cells in our experiments. We investigated impacts of cartridge volume, elution solvent, elution solvent volume, and sample pretreatment on halocode recoveries.

For all optimization experiments, 20uL of each halocoded nanoparticle formulation at a concentration of 1.5 mg/mL in milliQ water were pooled and spiked into an eppie with a cell pellet. 250 µL of RIPA lysis buffer with 0.9 w/v% SDS was added to each conical tube, vortexed, and incubated for 15 min at 65°C. The cell lysate was then sonicated for 2 seconds in a bath sonicator with a 1-minute rest period 3 times. Oasis HLB PRiME 3cc cartridges were used for SPE as the increased surface area can more easily accommodate increased viscosity of cell lysate.

For optimization of the elution solvent, spiked cell lysate samples were diluted at a 1:1 ratio with 4% phosphoric acid to acidify samples before SPE with a 20-position cartridge vacuum manifold. The SPE cartridge was conditioned with 2 mL of methanol and equilibrated with 1 mL of milliQ water. The sample was loaded at a flow rate of 1 mL/min, washed with 1 mL of water and 1 mL of 5% methanol, and then eluted with either 1mL of MeOH, 70ACN/30MeOH, 90ACN/10MeOH, or ACN at 1 mL/min. The halocode concentrations of eluted sample were determined with GC-MS and compared to the initial halocode concentration calculated based on the wt. % loading of the nanoparticle to determine the SPE efficiency for each halocode. We found that switching the elution solvent from methanol to acetonitrile, more effectively disrupts hydrophobic interactions between analyte and matrix, increasing the % recovery for every halocode.

For optimization of the elution solvent volume, we performed the same experiment but varied the volume of ACN used in the final elution step. We eluted with one elution of 1 mL of ACN, 2 elutions of 500uL ACN, and 3 elutions of 300uL ACN. We found that we were able to elute all halocodes from cartridge at this concentration with 2 elutions of 0.3, and thus adapted our protocol to concentrate our samples 3.3x as compared to yields from the initial protocol.

For optimization of the sample pretreatment, we compared the halocode recovery with the addition of 0% ACN, 5% ACN, and 10% ACN total organic solvent percentage after the acidification/dilution sample pretreatment. We found that adding 5% and 10% ACN to samples during sample acidification reduced halocode sequestration, likely from isoelectric protein precipitation.[9,10]

*SPE of Halocoded Nanoparticles with 3cc Oasis HLB PRiME Cartridge for Matrix Recovery Studies*

Halocoded nanoparticles at a concentration of 1.5 mg/mL in milliQ water were added to either water, PBS, full biological media used for cell culture (RPMI-1640 media + 10% FBS + 1% penstrep), or cell lysate. All nanoparticle samples were diluted at a 1:1 ratio with 4% phosphoric acid to acidify samples before SPE with a 20-position cartridge vacuum manifold. The nanoparticle samples in cell lysate were also simultaneously diluted with ACN to contain a total organic solvent concentration of 5% to prevent halocode entrapment by precipitated proteins during acidification. The sample was loaded at a flow rate of 1 mL/min, washed 1 mL of 5% methanol, and then eluted two times with 300µL of ACN at 1 mL/min. The halocode concentrations of eluted sample were determined with GC-MS/EC and compared to the initial halocode concentration calculated based on the wt. % loading of the nanoparticle to determine the SPE efficiency for each halocode.

*In vitro Dosing of Halocoded NP Library*

OVCAR8 cells were seeded at 200,000 cells/well in 0.75 mL RPMI-1640 media supplemented with 10% FBS in a 12-well plate. OVCAR8 cells were used up to passage 30. Cells were allowed to grow for 24 hours (37°C, 5% $CO_2$) prior to treatment with NPs. Prior to dosing, all halocoded PLGA nanoparticle formulations were normalized to a PLGA concentration of 1.3 mg/mL. Cells were separately dosed with 75 µL of normalized PLGA nanoparticles. The sides of the cell culture plate and lid were parafilmed to prevent evaporation, and cells and nanoparticles were incubated for 4 hours at 37°C and 5% $CO_2$.

After incubation, cells were washed once with 500µL of warm PBS and dissociated with 100µL 0.25% Trypsin-EDTA. Trypsin was quenched with 1mL of media and the cell suspension was pipetted up and down to ensure that all cells had been dissociated from the plate.

Cells were transferred to 1.7mL Eppendorf tubes and pelleted down at 300 rcf for 5 min. The supernatant was removed, and cells were resuspended in 50µL of media and counted. Tubes were parafilmed and stored at -20°C in preparation for extraction. For analysis, tubes corresponding to cells dosed with each library formulation were pooled in a conical tube. This corresponded to 5 total wells pooled per experiment.

250 µL of RIPA lysis buffer with 0.9 w/v% SDS were added to each conical tube, vortexed, and incubated for 15 min at 65°C. The cell lysate was then sonicated for 2 seconds in a bath sonicator with a 1-minute rest period 3 times. SPE was used to extract halocodes from cell lysate with the 3cc HLB PRiME cartridge and halocodes were detected with GC-MS using the procedures described above. % NP uptake was determined as a function of theoretical max of halocode dosed for each formulation and SPE % recovery for each halocode in cell lysate. The theoretical max dose was

determined from halocode wt. % loading. % NP uptake values were normalized to cell count.

*In vitro Dosing of Fluorescent NP Library*

OVCAR8 cells were seeded at 200,000 cells/well in 0.50 mL RPMI-1640 media supplemented with 10% FBS in a 12-well plate. OVCAR8 cells were used up to passage 30. Cells were allowed to grow for 24 hours prior to treatment with NPs. Prior to dosing, all halocoded PLGA nanoparticle formulations were normalized to a concentration of 0.5 mg/mL. Cells were dosed with 50 µL of normalized PLGA nanoparticles. The sides of the cell culture plate were parafilmed to prevent evaporation, and cells and nanoparticles were incubated for 4 hours at 37°C and 5% $CO_2$.

After incubation, cells were washed once with 500 µL of warm PBS and dissociated with 100µL 0.25% Trypsin-EDTA. Trypsin was quenched with 1mL of media and the cell suspension was pipetted up and down to ensure that all cells had been dissociated from the plate. Cells were transferred to a 1.7mL Eppendorf tube and pelleted at 300rcf for 5 min. The supernatant was removed, and cells were resuspended in 500µL of PBS and counted.

Nanoparticle-treated cells were aliquoted at 180 µL each into two wells of an opaque black 96-well plate. The fluorescence intensity was measured from the top with excitation and emission peaks of 646 and 676 nm with a BioTek Synergy H1 microplate reader. The acquisition settings including a gain of 120 remained the same for all experiments and creation of the standard curve. % NP uptake was determined as a function of the theoretical max of dosed fluorescent PLGA NPs. % NP uptake values were normalized to cell count and adjusted for autofluorescence measured with fluorescence from undosed control wells.

*Statistical Analysis*

Statistics were calculated using GraphPad Prism. Data are reported as mean ± standard deviation unless specified otherwise. Multiple comparisons were conducted using Kruskal-Wallis test with post hoc Dunn's multiple comparisons test. * $p < 0.05$, ** $p < 0.01$, *** $p < 0.001$.

*Deconvolution Microscopy*

Chambered cover glass was coated with rat tail collagen (300 µL of 50 µg/mL in 0.02N acetic acid). After 5 minutes, the wells were washed with room temperature PBS and allowed to dry in a sterile environment. Wells were stored at 4 °C up to one week prior to seeding OVCAR8 cells at 8,000 cells/well. The cells were allowed to adhere for 24 hours prior to treatment with 15 µL of a 0.5 mg/mL sulfoCy3 labeled-NP solution. The cells were incubated for 24 hours, at which time they were washed 3x with PBS and fixed in 4% methanol free formaldehyde prepared in PBS (15 min, room temperature). This was followed by three washes of ice-cold Hank's Balanced Salt Solution (HBSS, ThermoFisher) at 5 min/wash. Solutions and cover glass/wells were kept covered with aluminum foil for the following steps to prevent photobleaching of the fluorophores. 300 µL of wheat germ agglutinin-AF647 (10 µg/mL in HBSS) were added to each well and incubated at room temperature for 10 min. Cells were washed 3x with room temperature PBS (5 min/wash) before fixing again with 4% formaldehyde for 2 min followed by another

3x PBS wash (5 min/wash). 300 μL of 1.25 μg/mL Hoechst in PBS were added to each well for 2 min at room temperature before washing with 3x with room temperature PBS (5 min/wash). Finally, the wells were filled with ~5 drops of Vectashield (H1000) and stored at 4 °C protected from light. The cells were imaged with the Applied Precision DeltaVision Ultimate Focus Microscope with TIRF Module (Inverted Olympus X71 microscope) equipped with 405, 488, 512, and 568 nm lasers. Images were acquired with 60 and 100x objectives. All images were acquired under the same illumination settings and then processed with OMX softWoRx software (Applied Precison/GE). Image LUTs were linearly adjusted to improve contrast on FIJI. Z slices were merged into Z projections.

*Flow Cytometry*

OVCAR8 cells were seeded at 8,000-10,000 cells/well in 100 µL RPMI-1640 supplemented with FBS and pen/strep in a 96-well plate. Cells were allowed to grow for 24 hours prior to nanoparticle treatment. PLGA Cy5 nanoparticle formulations were normalized based on their fluorescence intensities, and cells were dosed with 5 µL of normalized particles. Cells and nanoparticles were incubated for 24 hours at 37°C. After incubation, cells were washed once with PBS and dissociated with 40 µL warmed 0.25% trypsin-EDTA for 5 minutes at 37°C. Trypsin was quenched with 180 µL cold media and the plate was placed on ice. Samples were analyzed using a BD LSR II Flow Cytometer with a high throughput sampler (BD Biosciences) on the APC channel (ex. 640, filters 670/30). Data were analyzed using FlowJo.

*Liposome Synthesis*

Liposomes comprising 20 mol% cholesterol, 36 mol% DSPC, 36 mol% DSPG, 8 mol% 18:0 propargyl PC were synthesized via thin film hydration, extruded to 50 nm, and characterized, as described previously.[1,11]

*Halocode Linker Synthesis*

Reaction conditions were adapted from Ohlmeyer et al.[12] Bromo-PEG1-azide (50 mg, 0.26 mmol) and respective chlorophenol (1.2 eq., 0.31 mmol) were dissolved in dimethylformamide (DMF, 1 mL). Cesium carbonate (100.8 mg, 0.31 mmol) were added, and the reaction was stirred at 80°C for two hours under $N_2$. Toluene (2.5 mL) was added, and the organic phase was washed twice with 2.5 mL 0.5 M NaOH, twice with 2.5 mL 1M HCl, and once with 2.5 mL water. The organic layer was dried over $MgSO_4$ and solvent was removed via rotary evaporation to yield a yellow oil. $^1H$ NMR and $^{13}C$ NMR were performed on a Bruker Avance-III HD Nanobay spectrometer operating at 400.09 or 400.13 MHz.

*Halocoding of Liposomes via Copper Catalyzed Azide-Alkyne Cycloaddition*

Liposomes were covalently halocoded via copper catalyzed azide-alkyne cycloaddition as described previously.[11] Briefly, liposomes (2 mg, 1 mg/mL, in water) were mixed with 3-chlorophenol-EG-$N_3$, 3,5-dichlorophenol-EG-$N_3$, or 2,3,6-trichlorophenol-EG-$N_3$ at 10 molar eq. of azide linker with respect to 18:0 propargyl PC lipid, 5.9 mL sodium ascorbate (1767 μM final concentration), and 5.9 mL $CuSO_4$/THPTA solution (50 μM $CuSO_4$, 100 μM THPTA final concentration). All reactions were carried out at room temperature protected from light over 12-16 hours. Subsequently, TFF was used to remove excess reagents.

*MALDI Tissue Preparation*

Approximately 5 g of untreated control mouse brain was removed from -80°C storage, homogenized using a gentleMACS Dissociator, and transferred to a 3x3 cm plastic mold for freezing. The resulting frozen brain homogenate was sectioned onto an indium-tin-oxide (ITO) (Catalog: 703192, Sigma Aldrich) slide compatible for MALDI MSI. Varying concentrations (150, 100, 50, 25, 10 μM) of 3-chlorophenol-EG-$N_3$, 3,5-dichlorophenol-EG-$N_3$, and 2,3,6-trichlorophenol-EG-$N_3$ were spotted on top of brain homogenate at a volume of 1 μL and placed in a desiccator. The protocol for spotting standards onto control tissue for quantitative analysis was adapted from previous publications.[13,14] Once dry, spotted brain homogenate sections were coated with 2,5-dihydroxybenzoic acid (DHB, 40 mg/mL) in 50:50 methanol: water with 0.01% trifluoroacetic acid (TFA) applied directly onto the ITO slide using a robotic sprayer (TM-sprayer, HTX imaging, Carrboro, NC) with a twelve-pass cycle, flow rate (0.05 mL/min), spray nozzle heat (75 °C), track spacing (3 mm), nozzle velocity (1200 mm/min), and 10 psi nitrogen pressure.

Gelatin tissue microarray (TMA) mimetic molds were prepared at 40% concentration. Molds consisted of 1.5mm core diameter wells (Ted Pella Inc., Redding, CA). Homogenized control mouse liver was prepared using the gentleMACS Dissociator and aliquoted into 15 microcentrifuge tubes. Aliquoted liver homogenate was then spiked with varying concentrations of 3-chlorophenol-EG- $N_3$, 3,5-dichlorophenol-EG-$N_3$, and 2,3,6-trichlorophenol-EG-$N_3$, from 10 μM to 150 μM. Mimetic tissue model protocol was adapted from previous publications.[15] Each nanoparticle formulation had its own respective row for individual quantitation. Spiked tissue standards were frozen at -80 °C and transferred into the TMA gelatin mold using a 1.5mm core punch tool (Ted Pella Inc., Redding, CA). Mimetic molds were stored at -80 °C until analysis. Frozen molds were sectioned at 10 μm thickness and thaw mounted directly onto ITO slides. DHB matrix was applied to the MALDI MSI mimetic sections following the TM sprayer protocol described above.

*MALDI Imaging*

MALDI mass spectrometry images were acquired in positive ion mode using a timsTOF flex mass spectrometer (Bruker LLC, Billerica, MA). Mass range for the acquisition was set from *m/z* 200-2000. 3-chlorophenol-EG- $N_3$, 3,5-dichlorophenol-EG-$N_3$, and 2,3,6-trichlorophenol-EG- $N_3$ formulations were detected using full scan mode to monitor *m/z* 1055.69, 1089.65, and 1123.61, respectively. Ion images were acquired with a pixel size of 100 μm with 800 shots per pixel and a laser frequency of 10,000 Hz. A global mass calibration was performed prior to acquisition using electrospray ionization (ESI) source with a tune mix solution (Agilent Technologies, Santa Clara, CA). Identification of nanoparticle peaks were based on accurate mass. Ion images were visualized in SCILS Lab Software (Version 2022a, Bruker LLC, Billerica, MA).

<u>**Supplemental Figures**</u>

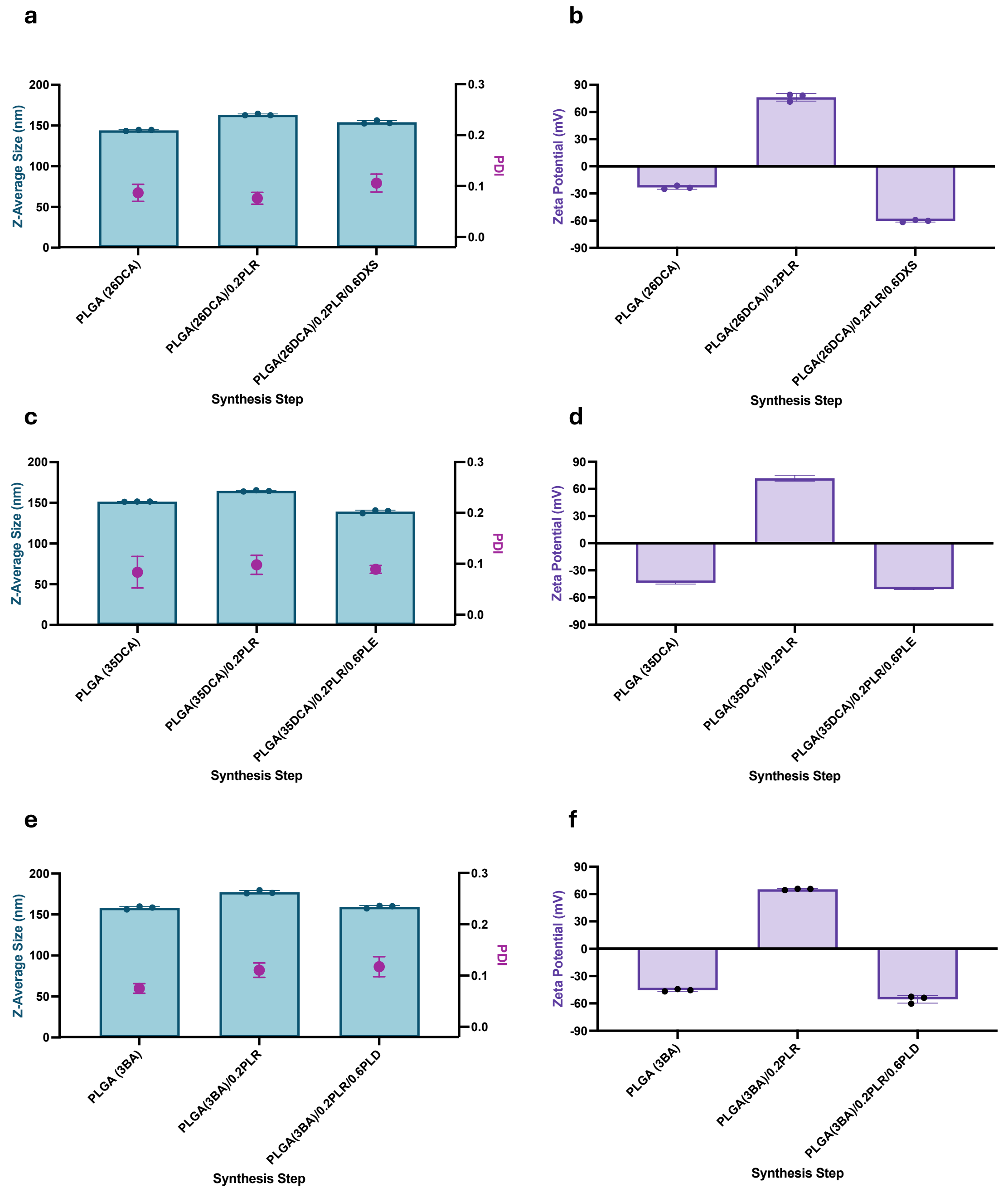


**Figure S1. DLS characterization of surface-modified halocoded library formulations during LbL synthesis process.** Anionic PLGA NPs were layered with 0.2 wt. eq. of PLR followed by 0.6 wt. eq. of each anionic polyelectrolyte. (a) Hydrodynamic diameter and PDI of PLGA-DXS halocoded NPs during each layering step of LbL surface modification measured by DLS. (b) Zeta potential in milliQ $H_2O$ of PLGA-DXS halocoded NPs during each layering step of LbL surface modification measured by electrophoretic light scattering. (c) Hydrodynamic size and PDI of PLGA-PLE halocoded NPs during each layering step of LbL surface modification measured by DLS. (d) Zeta potential in milliQ $H_2O$ of PLGA-PLE halocoded NPs during each layering step of LbL surface modification measured by electrophoretic light scattering. (e) Hydrodynamic size and PDI of PLGA-PLD halocoded NPs during each layering step of LbL surface modification measured by DLS. (f) Zeta potential in milliQ $H_2O$ of PLGA-PLD halocoded NPs during each layering step of LbL surface modification measured by electrophoretic light scattering. Data shown as mean ± SD (n=3).

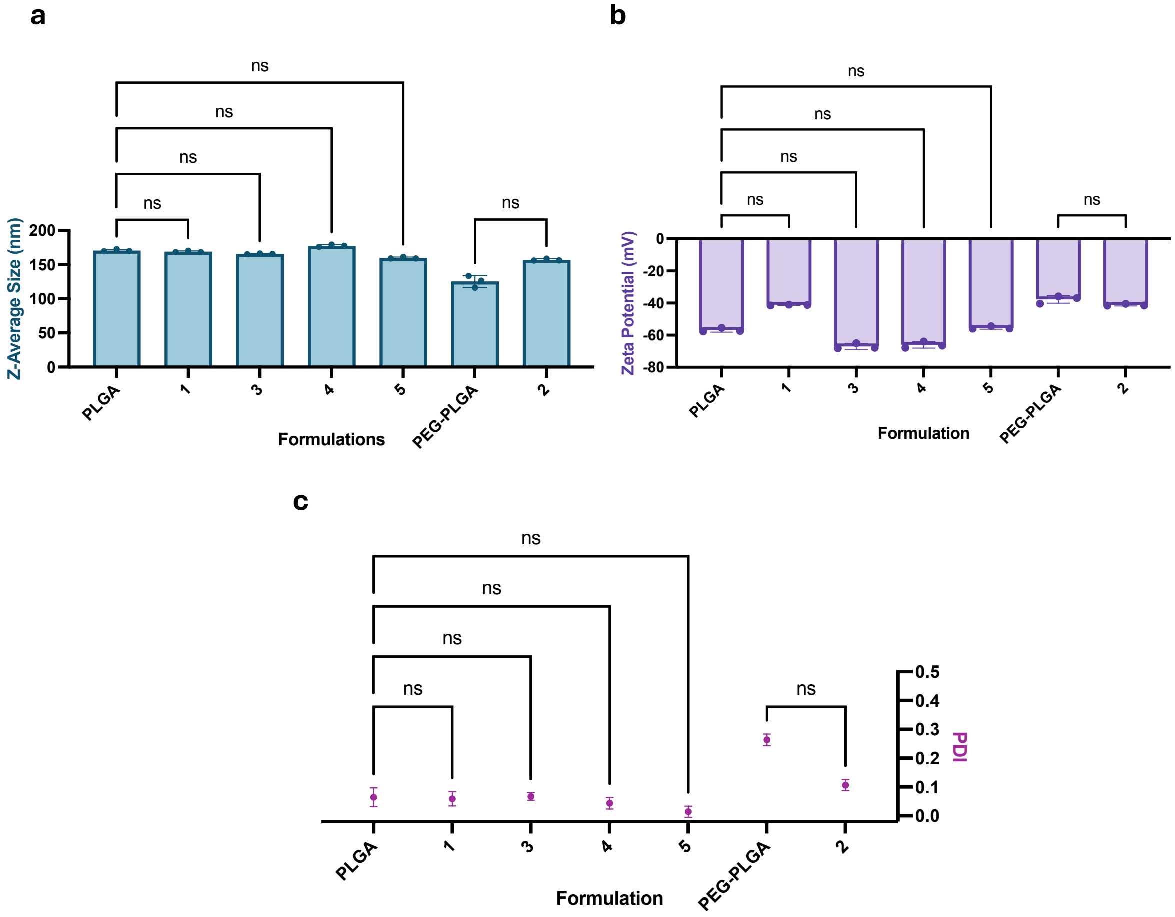


**Figure S2. Comparison of physicochemical properties of bare PLGA and PEG-PLGA NPs to halocoded library formulations.** There was no significant change to (a) hydrodynamic size (b) zeta potential or (c) PDI of NP formulations upon encapsulation of halocodes. Data shown as mean ± SD (n=3).

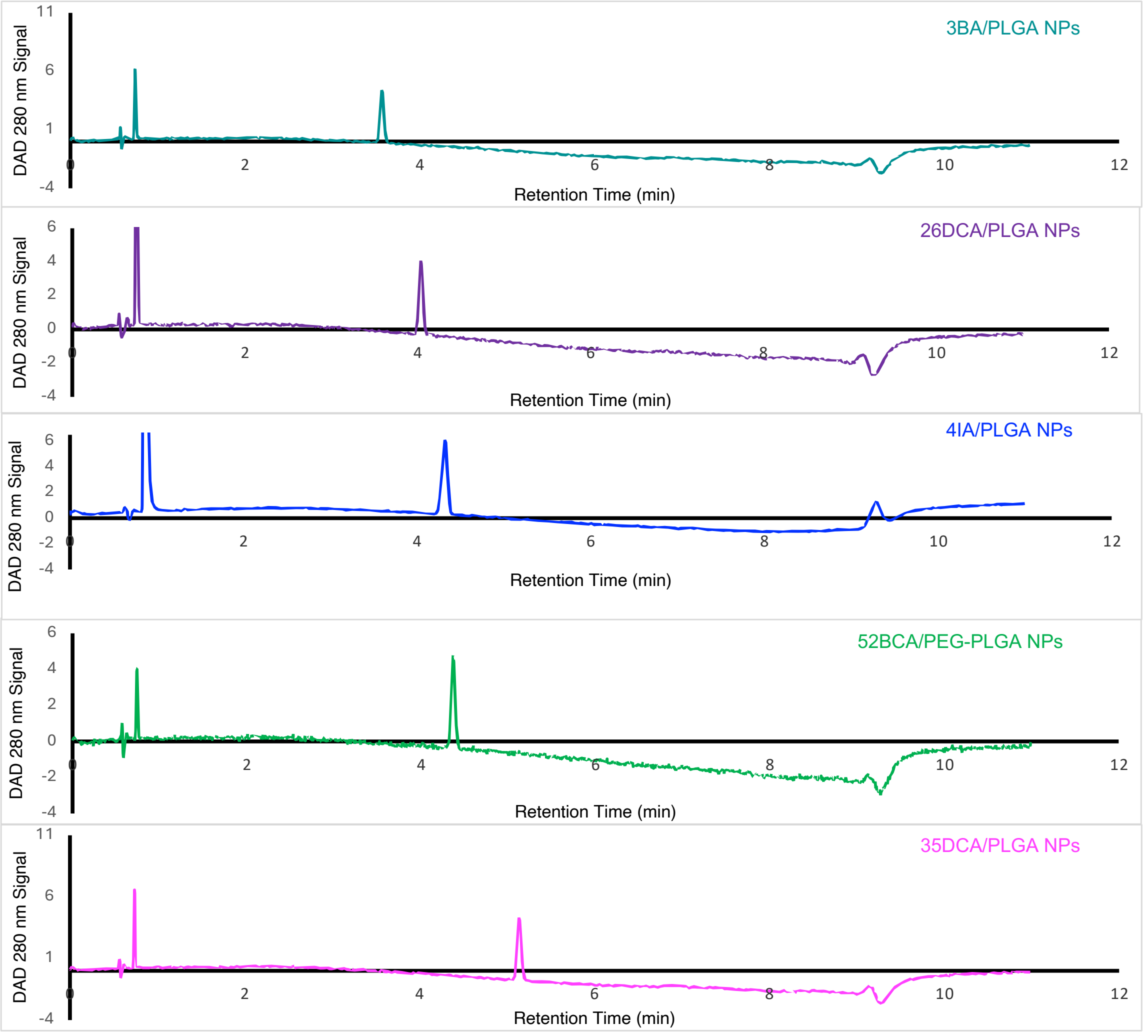


**Figure S3. Halocode loading of PLGA NP library.** HPLC traces with UV halocode detection for determining encapsulation efficiency and wt. % loading of the halocoded nanoparticle library. Halocodes elute in order of increasing polarity.

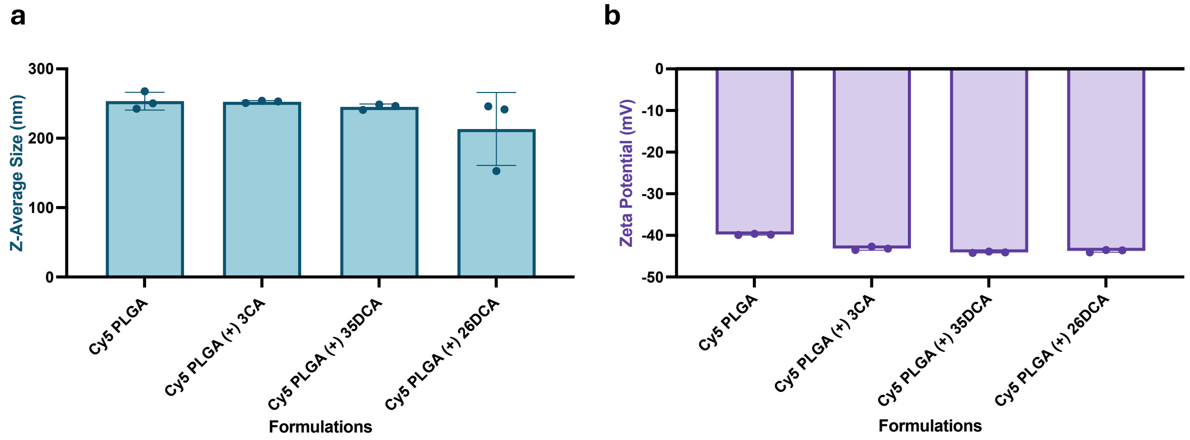


**Figure S4. Characterization of PLGA NPs coloaded with halocodes at 3 wt.% and fluorescent dye at 1 wt. %.** (a) Hydrodynamic Size of PLGA/Cy5/Halocode NPs measured by DLS. (b) Zeta potential of PLGA/Cy5/Halocode NPs in milliQ $H_2O$ measured by electrophoretic light scattering. Data shown as mean ± SD (n=3).

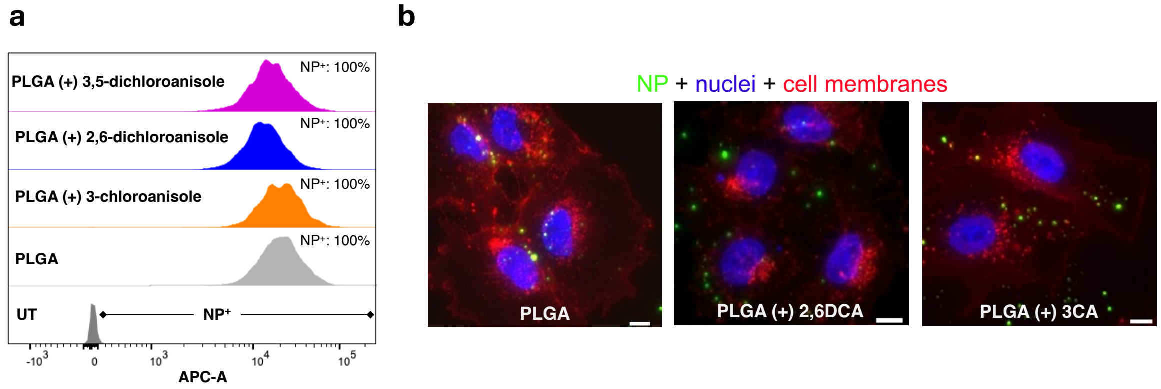


**Figure S5. Comparing delivery properties of halocoded NPs.** (a) Nanoparticle-cell association and (b) representative cell images of fluorescent halocoded PLGA NPs compared to fluorescent PLGA NPs 24 hours after dosing in an OVCAR8 cell line. Cell nuclei were stained with Hoechst3342 (blue), cell membranes with wheat germ agglutinin (red), and nanoparticles have encapsulated Cy5 dye (green). Scale bar = 10 µm. We elected to use a higher loading of halocodes (3 wt.%) in this study as disruption of nanoparticle properties would more easily occur at higher aryl halide loading levels. Both nanoparticle-cell association and nanoparticle localization remained similar for halocoded and fluorescent PLGA NPs, indicating that halocodes have minimal effects on delivery properties of NPs.

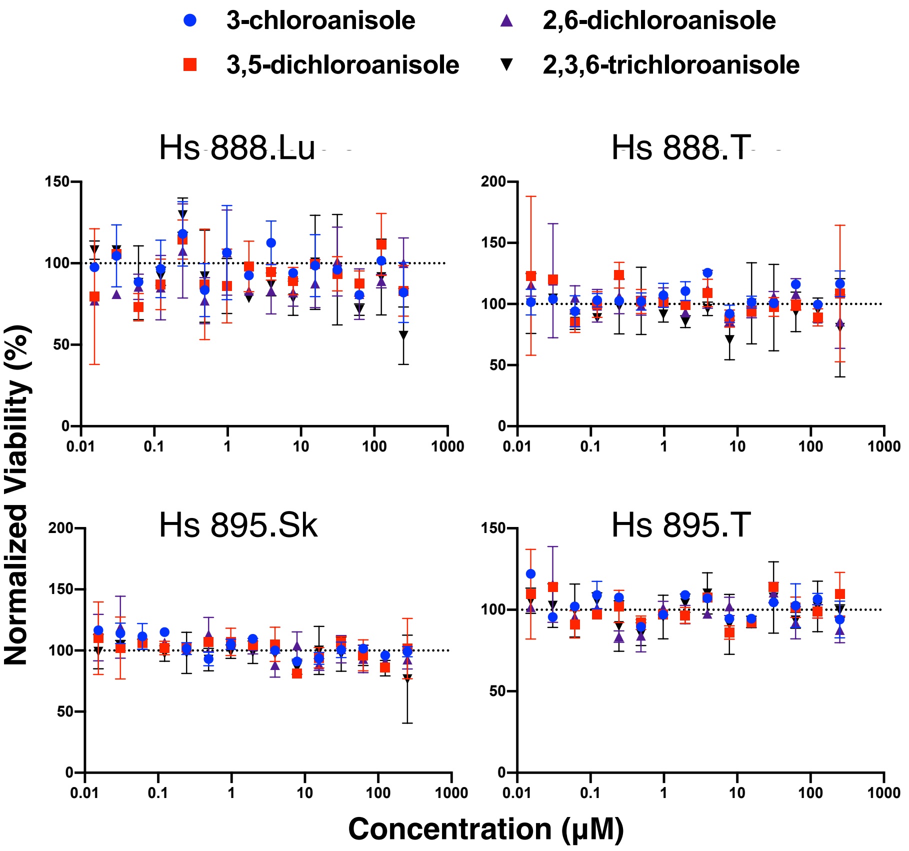


**Figure S6. Halocode biocompatibility for in vitro studies**. Cell viability 72 hours after representative tri and dichlorinated haloanisole dosing to (a) 888T, (b) 888Lu, (c) 895T, and (d) 895sk human normal and cancer cell lines was determined with CellTiter-Glo Luminescent Cell Viability Assay. Results were normalized to vehicle (DMSO) controls. n =2, individual data points shown.

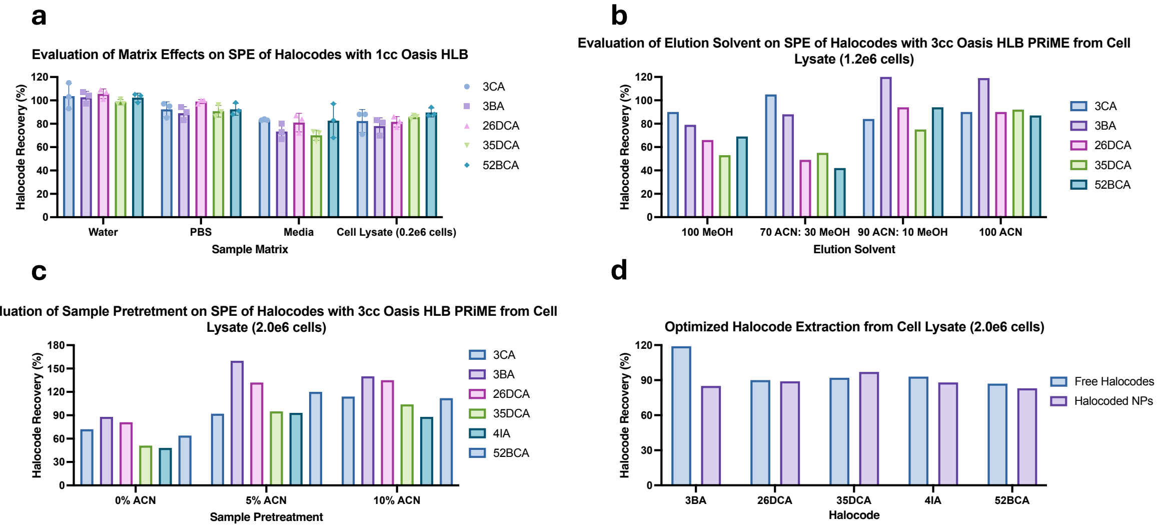


**Figure S7. SPE optimization of halocodes from biological matrices.** (a) Halocode recovery from different matrices using base protocol from Waters with 1cc Oasis HLB

SPE cartridges. The base protocol is described in detail in the methods section. Data are shown as mean ± SD (n=3). (b) Optimizing the choice of elution solvent when extracting halocodes spiked into a dense cell lysate matrix of 1.2e6 cells using 3cc Oasis HLB PRiME SPE cartridges. (n=1). (c) Optimizing the sample pretreatment when extraction halocodes from dense cell lysate matrix of 2.0e6 cells using 3cc Oasis HLB PRiME SPE cartridges and 100% ACN elution. (n=1) (d) Halocode % recovery for spiked halocodes and halocoded NPs from a dense cell lysate matrix using our optimized SPE procedure with 3cc Oasis HLB PRiME SPE cartridges. Details on the complete procedure can be found in the methods section (n=1). We were confident in our interpretation of the trends in the SPE optimization experiments, so we proceeded without replicates.

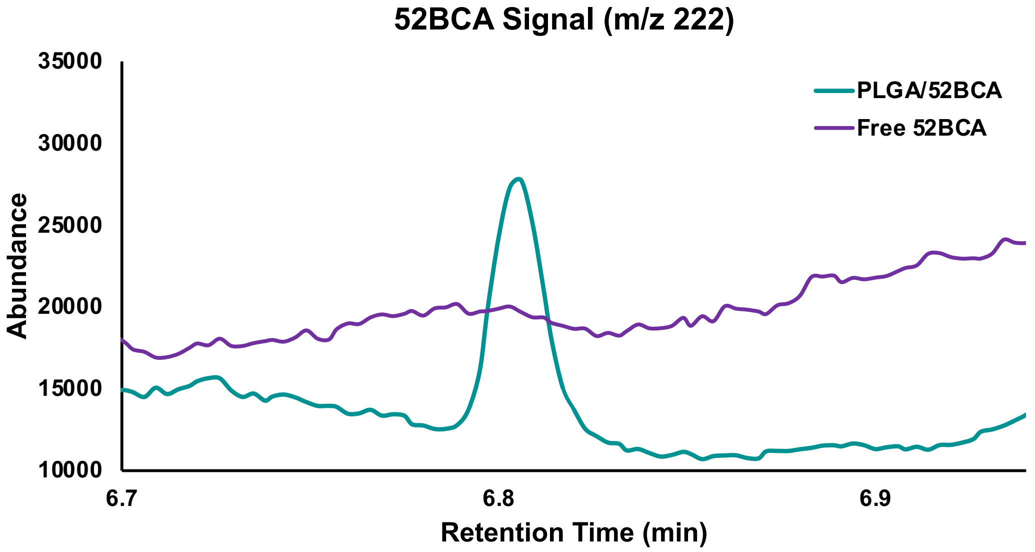


**Figure S8. Comparison of in vitro NP-cell association between encapsulated and free 52BCA** Overlaid representative chromatograms showing detection of the quantification ion (m/z 222) for 52BCA encapsulated inside of PLGA NPs versus freely dosed to OVCAR8 cells at the same concentration. There was no detectable peak for free 52BCA after dosing, incubation for 4 hours, and extraction from OVCAR8 cells while 52BCA was detectable for all wells dosed with NPs.

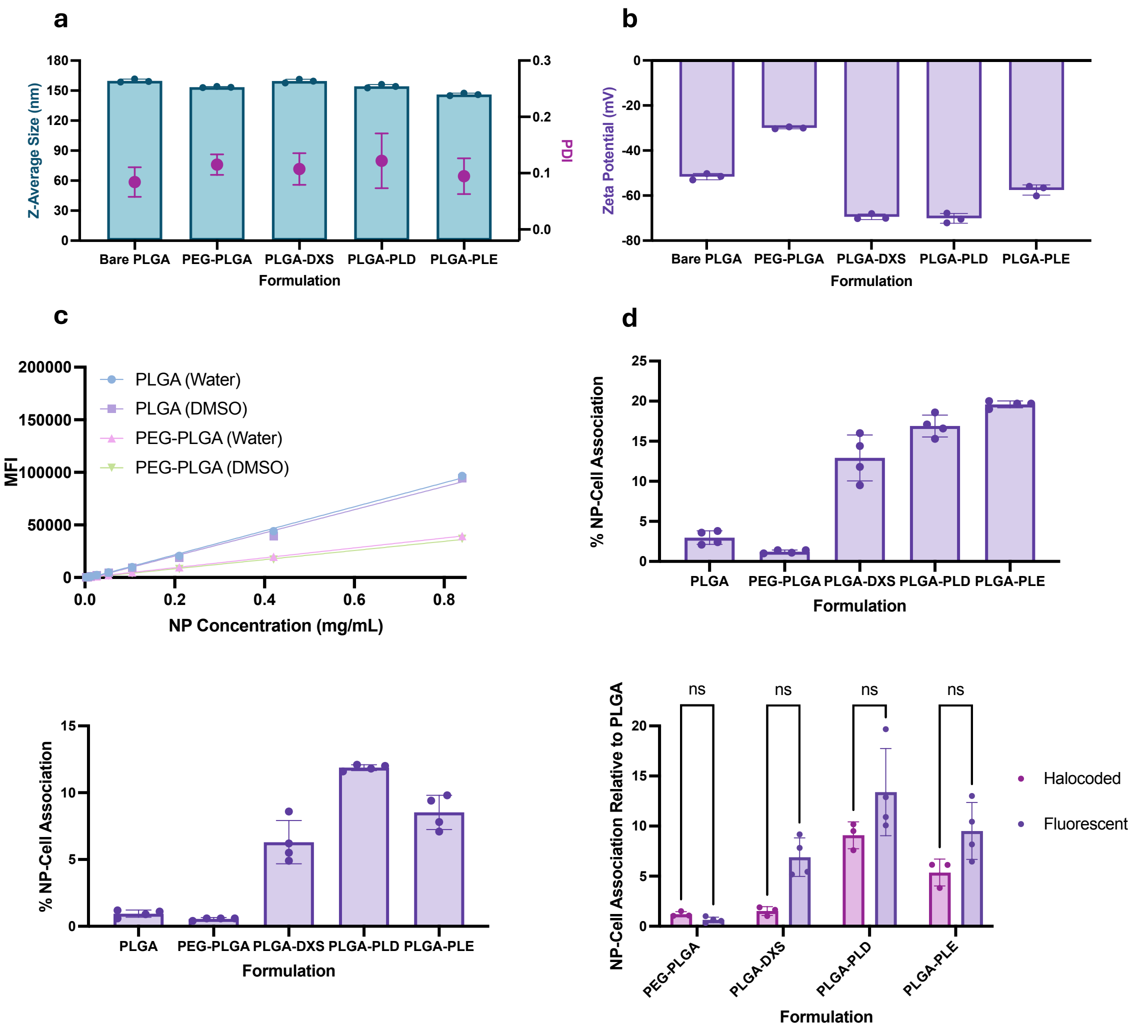


**Figure S9. Fluorescent NP Library Characterization.** (a) Hydrodynamic Size and PDI of Cy5 NP library by DLS. PLGA-Cy5 NPs were layered with 0.3 wt. eq of PLR followed by 1 wt. eq. of DXS, 1 wt. eq. of PLD, and 1.3 wt. eq. of PLE. (b) Zeta potential of Cy5 NP library in milliQ $H_2O$ measured by electrophoretic light scattering. Data shown as mean ± SD (n=3). (c) Standard curves showing fluorescence intensity for PLGA-Cy5 and PEG-PLGA-Cy5 formulations in measured with microplate spectrophotometry. Fluorescent measurements were taken in water and water + 25% DMSO. All curves have $R^2 > 0.99$. (d) NP-cell association of Cy5 NP library as a function of administered NP dose measured with cells suspended in water. (e) NP-cell association of Cy5 library as a function of administered NP dose with cells suspended in water + 25% DMSO. (f) NP-cell association of fluorescent NP library relative to bare PLGA in water + 25% DMSO (n=4) compared to halocoded library (n=3). Data are shown as mean ± SD.

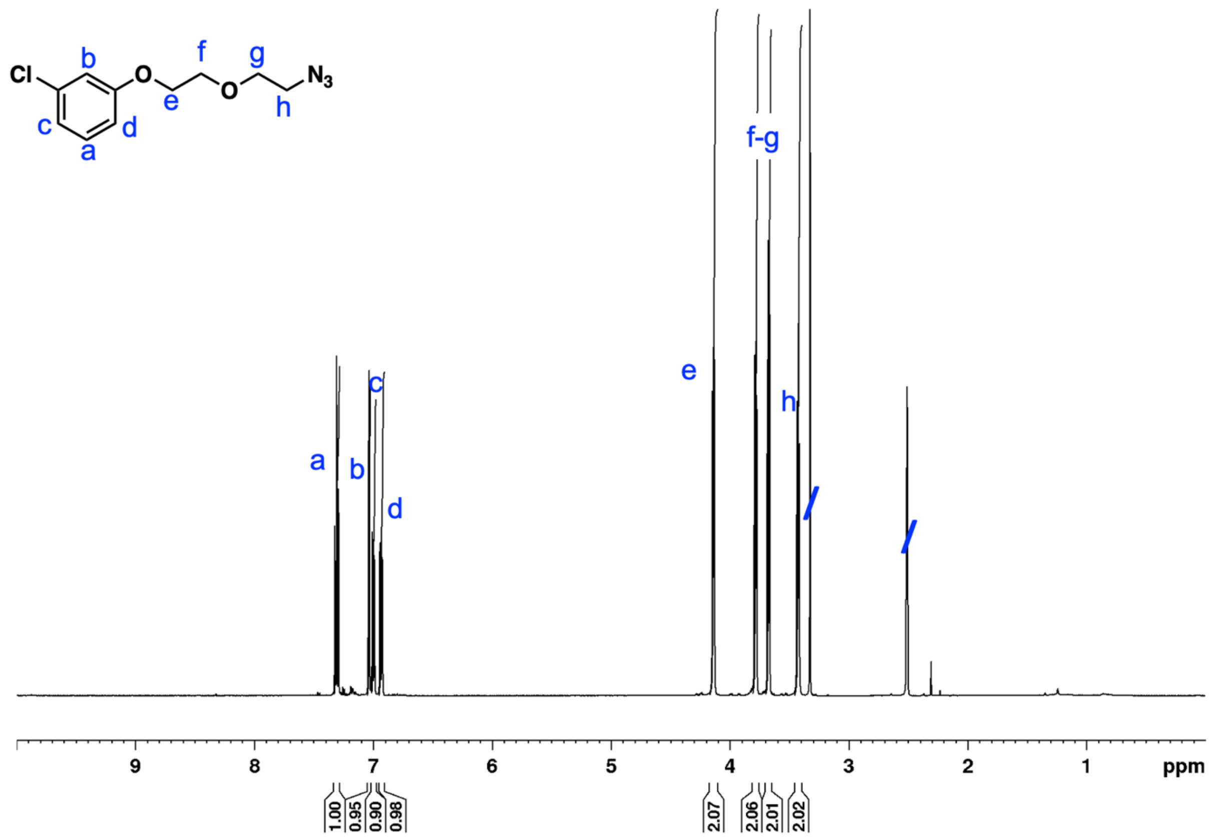

Figure S10. $^1$H NMR spectrum of 3-chlorophenol-EG-$N_3$, acquired in DMSO-$d_6$.

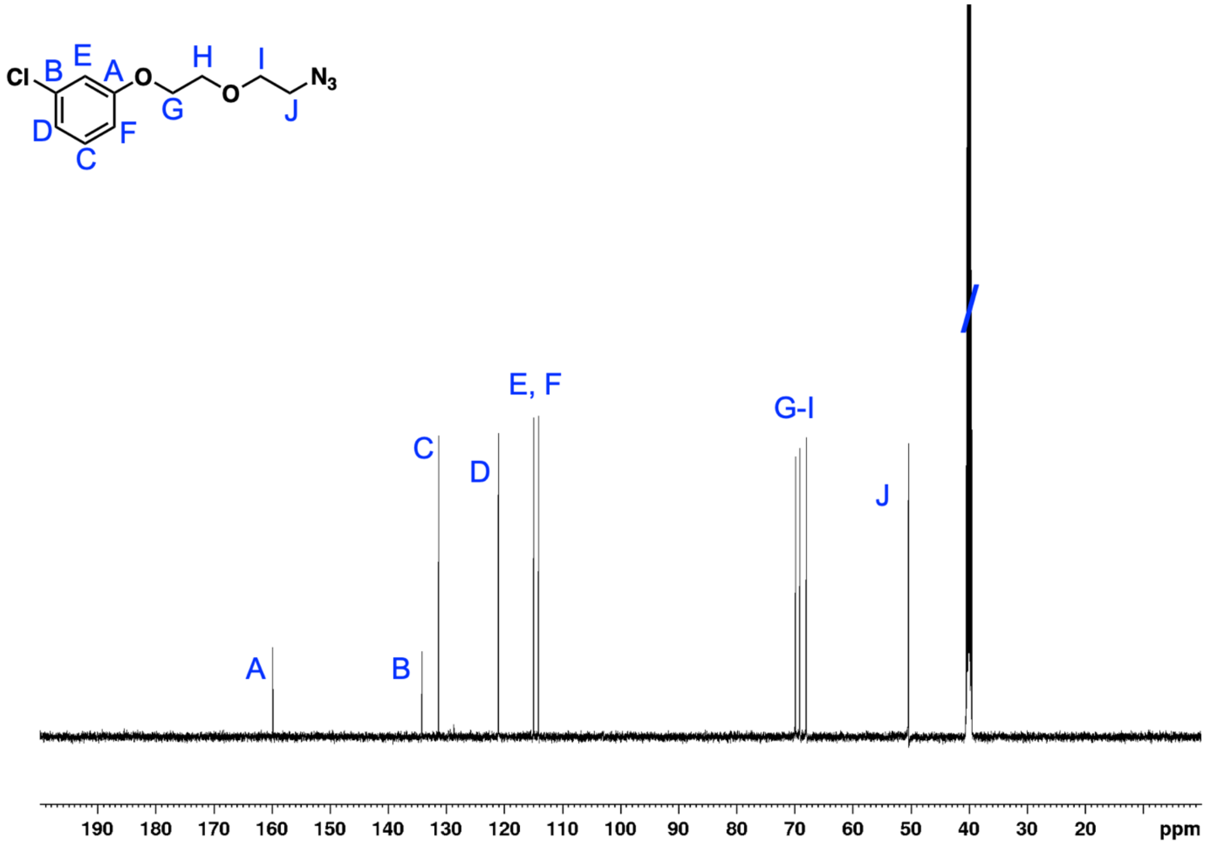


Figure S11. $^{13}C$ NMR spectrum of 3-chlorophenol-EG-$N_3$, acquired in DMSO-$d_6$.

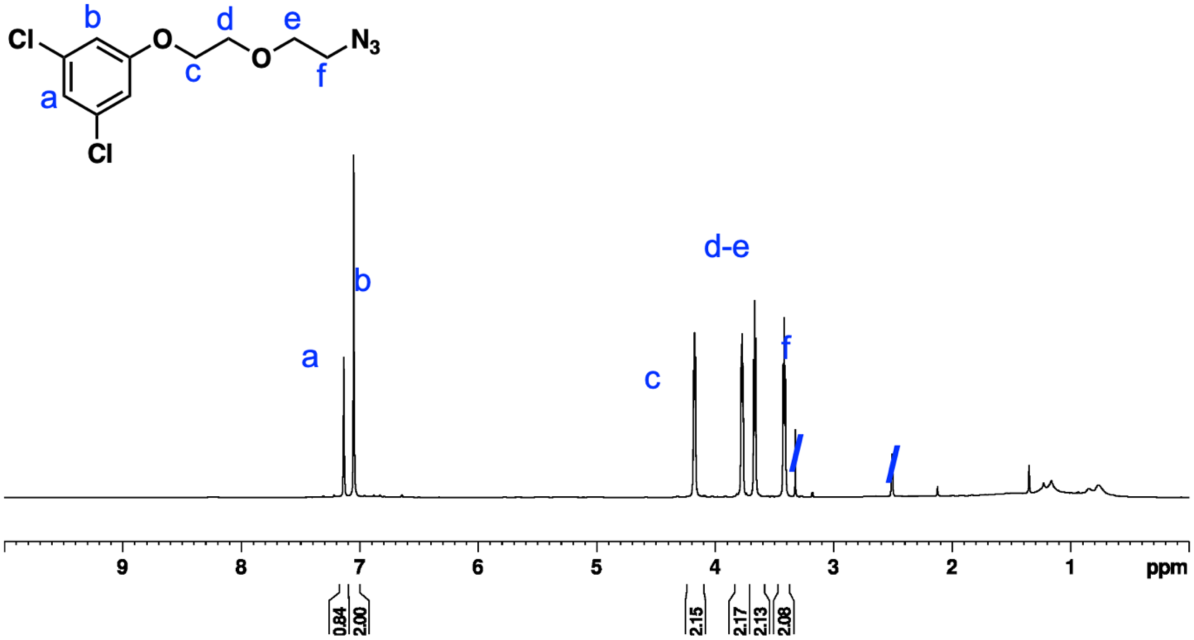


Figure S12. $^1H$ NMR spectrum of 3,5-dichlorophenol-EG-$N_3$, acquired in DMSO-$d_6$.

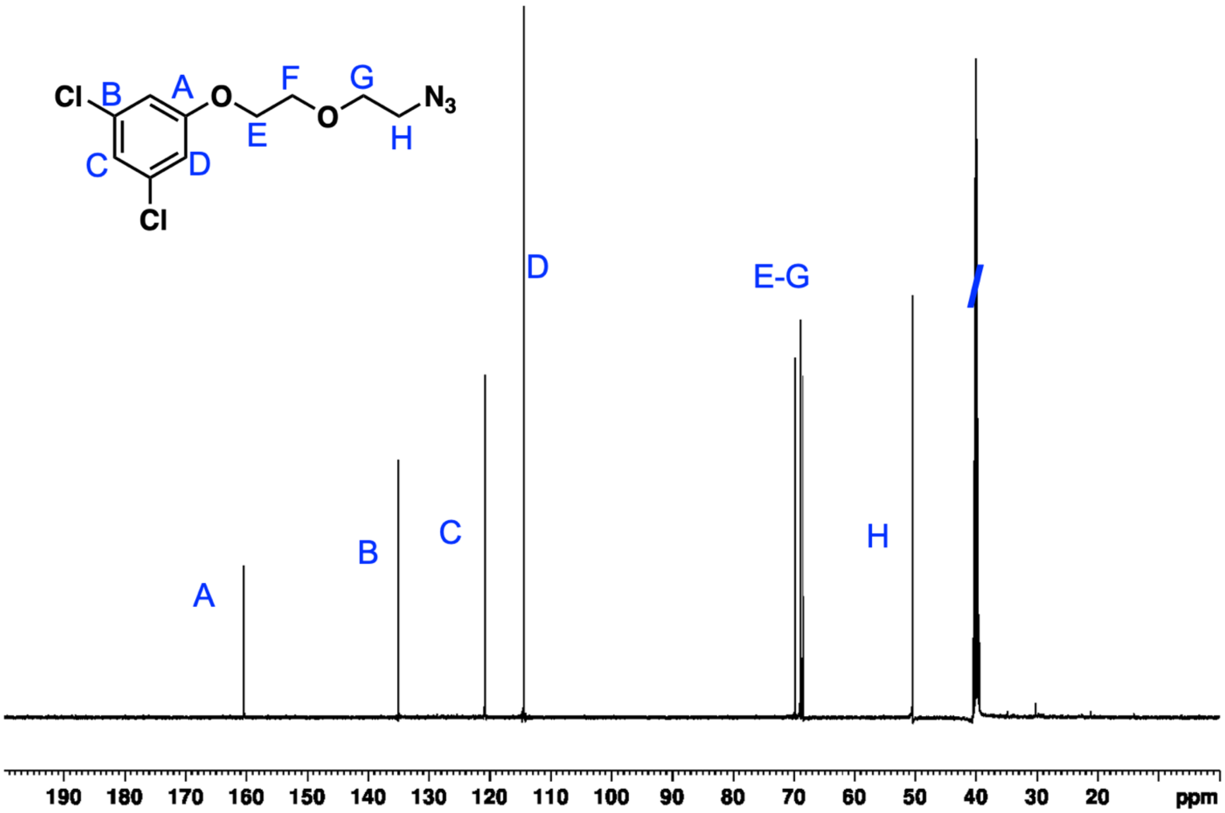


Figure S13. $^{13}C$ NMR spectrum of 3,5-dichlorophenol-EG-$N_3$, acquired in DMSO-$d_6$.

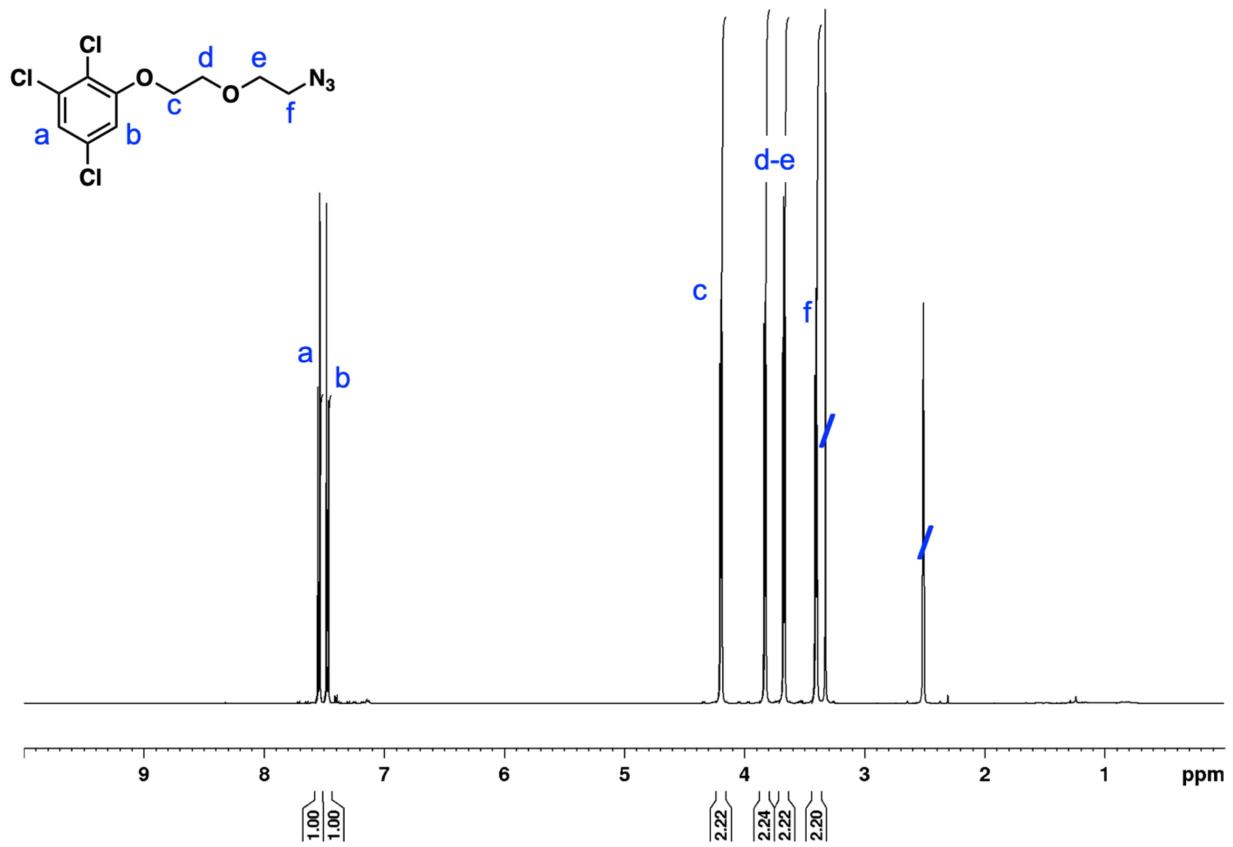


Figure S14. $^1$H NMR spectrum of 2,3,6-trichlorophenol-EG-$N_3$, acquired in DMSO-$d_6$.

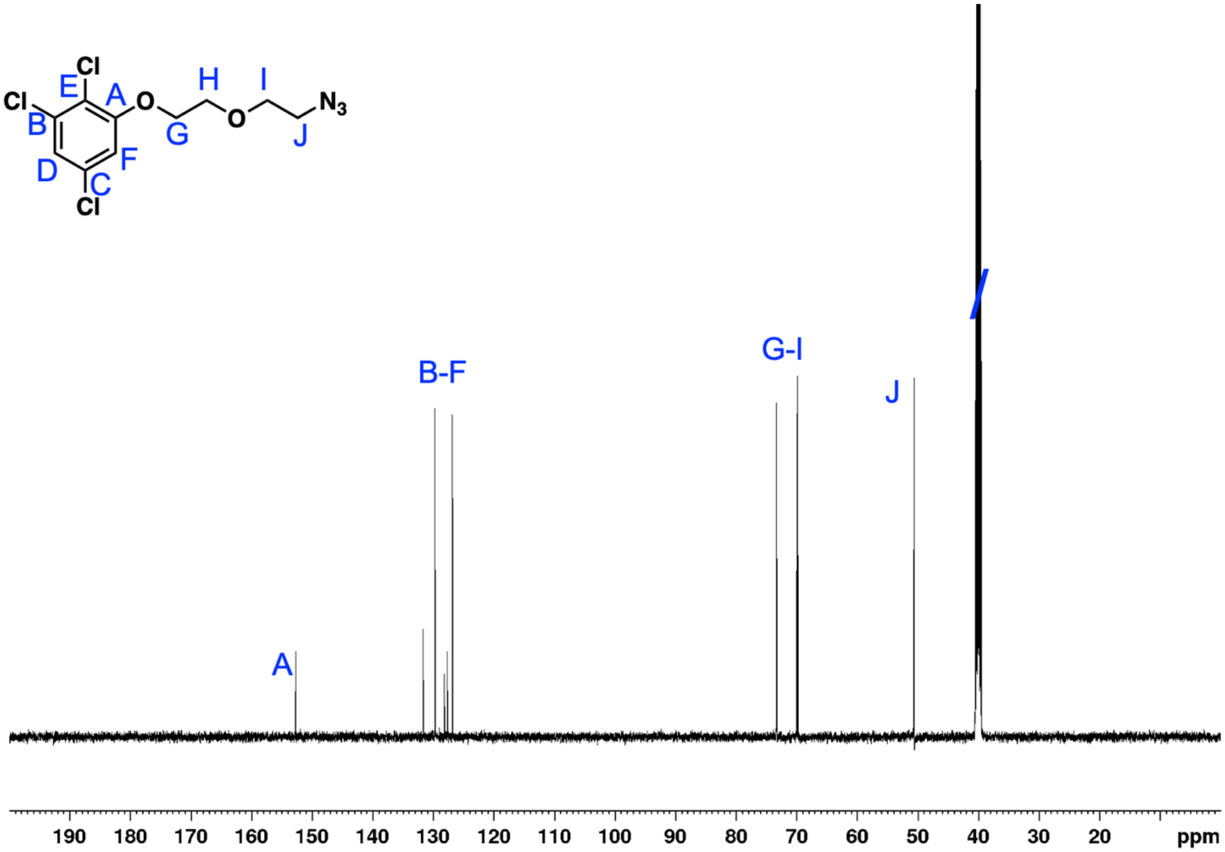


Figure S15. $^{13}C$ NMR spectrum of 2,3,6-trichlorophenol-EG-$N_3$, acquired in DMSO-$d_6$.

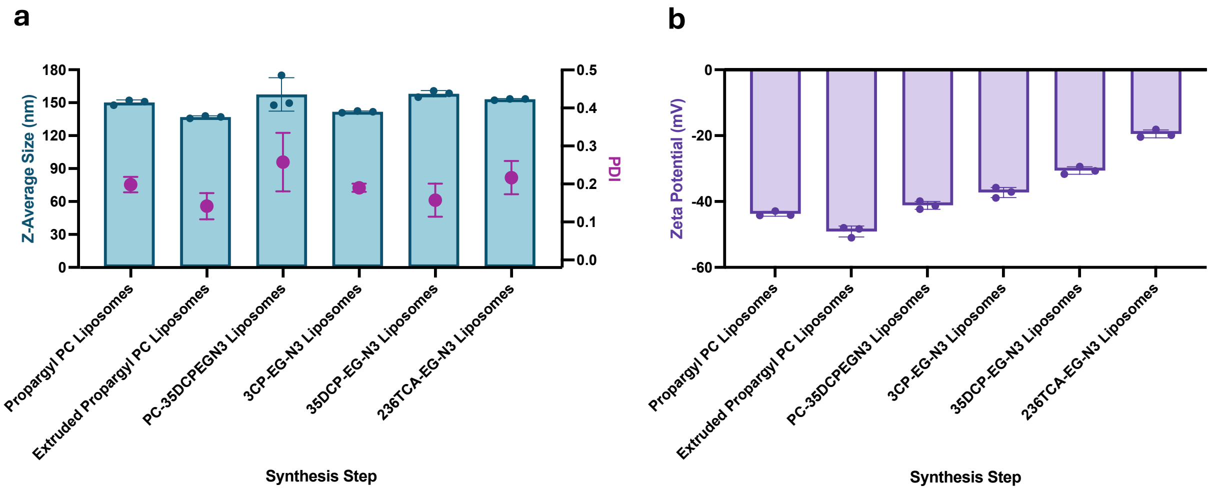


**Figure S16. Characterization of liposomes pre- and post-halocoding**. (a) Hydrodynamic Size of liposomes during the halocoding process measured by DLS. (b) Zeta potential of liposomes during the halocoding process in milliQ $H_2O$ measured by electrophoretic light scattering. Data shown as mean ± SD (n=3).

a

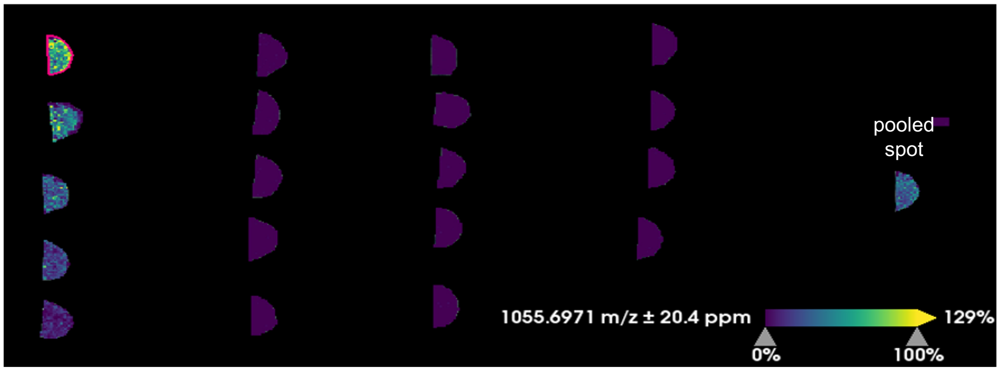


b

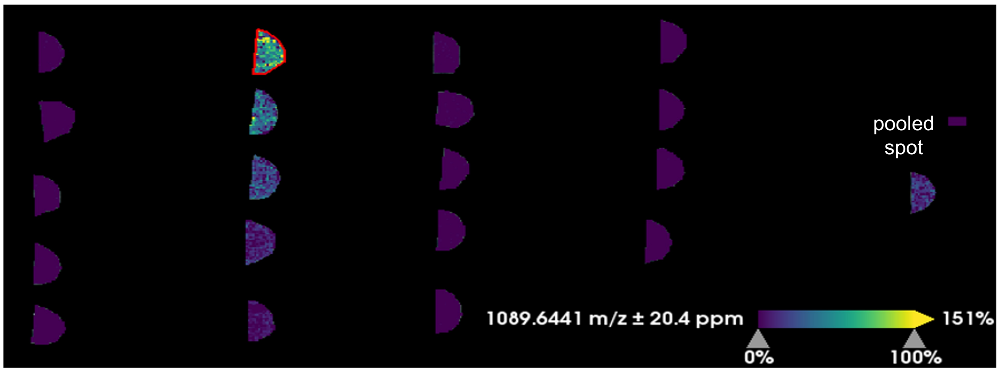


c

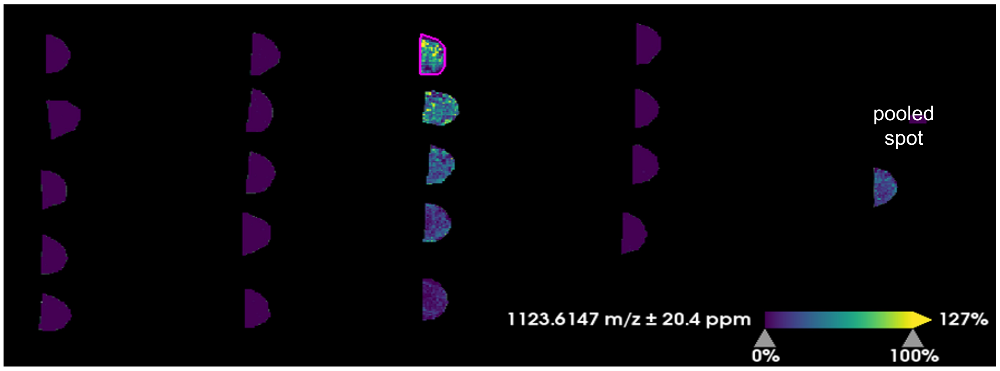


**Figure S17. MS signal intensities of halocoded lipids spotted onto tissue homogenate.** Integrated signal intensities corresponding to *m/z* values of (a) 1055.7 (3CP-EG-lipid); (b) 1089.6 (35DCP-EG-lipid); (c) 1123.6 (236TCP-EG-lipid) are shown. The pooled spot refers to a pooled mixture of the three halocoded liposomes which was deposited onto the tissue mimetic.

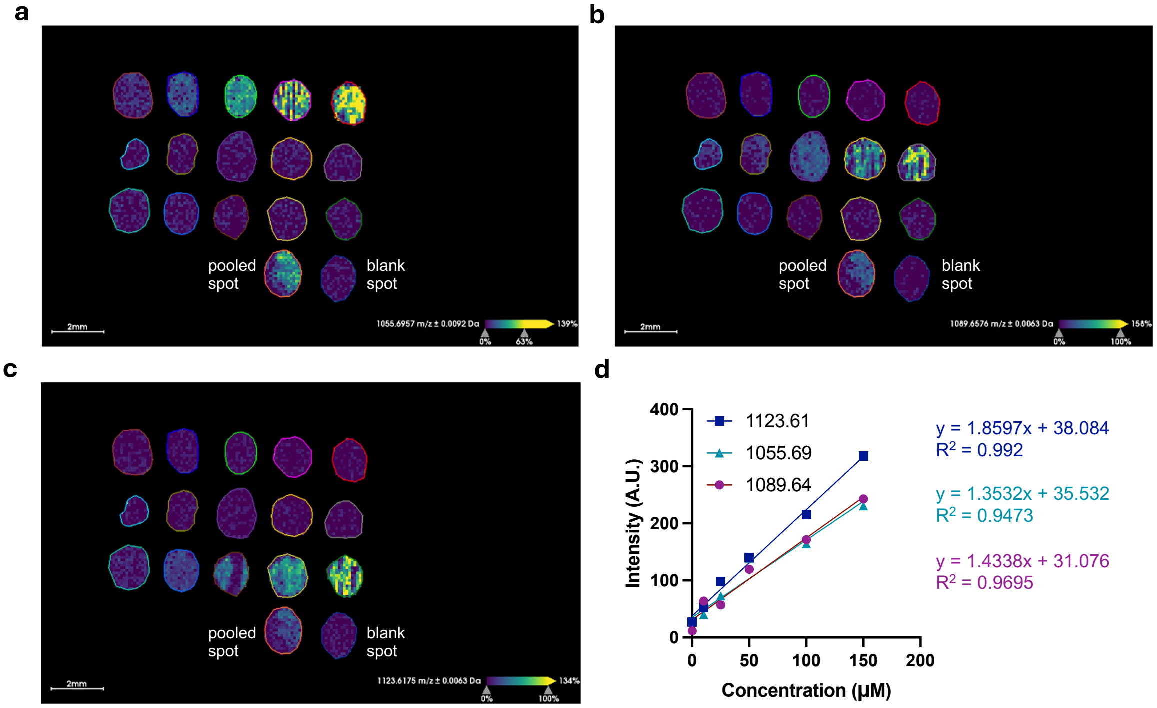


**Figure S18. MS signal intensities of halocoded lipids incorporated into liver mimetic**. Integrated signal intensities corresponding to *m/z* values of (a) 1055.7 (3CP-EG-lipid); (b) 1089.6 (35DCP-EG-lipid); (c) 1123.6 (236TCP-EG-lipid) are shown. (d) Expanded information for signal-response curves of integrated halocoded lipid signal intensities.

### Supplemental Tables

**Table S1. Linear halocode detection with MSD. LOD (S/N=3), LOQ (S/N=10)**

| Halocode | RT (min) | MSD Linear Fit $R^2$ | LOD (nM) | LOQ (nM) |
|---|---|---|---|---|
| 3BA | 5.9 | 1.0 | 0.74 | 2.5 |
| 26DCA | 5.9 | 1.0 | 0.60 | 2.0 |
| 35DCA | 6.2 | 1.0 | 0.78 | 2.6 |
| 4IA | 6.4 | 1.0 | 0.62 | 2.1 |
| 52BCA | 6.7 | 1.0 | 0.78 | 2.6 |

**Table S2.** Loading and encapsulation efficiency for the halocoded NP library. Data shown as mean ± SD (n=3).

| Formulation | Core Material | Halocode | Weight Percent Loading of Halocode (%) | Encapsulation Efficiency (%) |
|---|---|---|---|---|
| 1 | PLGA | 4IA | 0.58±0.01 | 5.8±0.1 |
| 2 | PEG-PLGA | 52BCA | 0.60±0.01 | 2.4±0.1 |
| 3 | PLGA | 26DCA | 0.64±0.06 | 2.6±0.2 |
| 4 | PLGA | 3BA | 0.63±0.03 | 2.5±0.1 |
| 5 | PLGA | 35DCA | 0.51±0.01 | 2.0±0.1 |
